# MAC3A and MAC3B modulate *FLM* splicing to repress photoperiod-dependent floral transition

**DOI:** 10.1101/2024.03.26.586198

**Authors:** Yu-Wen Huang, Chih-Yen Tseng, Yi-Tsung Tu, Hsin-Yu Hsieh, Yu-Sen Wang, Yun-Tung Ly, Yu-Zhen Chen, Shih-Long Tu, Chin-Mei Lee

## Abstract

Plants adjust their flowering time by integrating environmental cues through complex regulatory networks. RNA splicing plays a crucial role in modulating gene expression in response to flowering signals. The MOS4-associated complex (MAC), consisting of the evolutionarily conserved E3 ubiquitin ligases MAC3A and MAC3B, is pivotal in splicing regulation. However, their involvement in floral transition remained unclear. This study observed that *mac3a/mac3b* mutants flowered significantly earlier under short-day (SD) conditions, a phenotype absent under long-day (LD) conditions. This early flowering correlated with upregulation of *FLOWERING LOCUS T* (*FT*) and *SUPPRESSOR OF OVEREXPRESSION OF CO 1* (*SOC1*) compared to wild-type plants. Transcriptomic analysis revealed alterations in transcript levels and splicing profiles of key floral regulators across different flowering pathways. Further investigation identified the thermosensory flowering regulator *FLOWERING LOCUS M* (*FLM*) as being influenced by *MAC3A* and *MAC3B*. Subsequently, we found that *MAC3A* and *MAC3B* exhibited higher expression and were associated with *FLM* transcripts to modulate their splicing in SD. This study elucidates how the MAC complex, through RNA splicing regulation, integrates environmental signals to modulate flowering, unveiling a new layer of complexity in flowering pathways crosstalk under non-inductive photoperiods.

## Introduction

The floral transition from vegetative to reproductive stages represents a captivating and distinctive developmental process in angiosperms. To ensure successful reproduction, plants have evolved a sophisticated network that integrates various environmental signals, managing the plants’ developmental and metabolic status to precisely control floral transition through multifaceted mechanisms (Song et al., 2015; Park et al., 2019; Gendron et al., 2021; Takagi et al., 2023). This intricate regulatory network is categorized into major pathways, including photoperiod, age, vernalization, autonomous, thermosensory, and gibberellin (GA) pathways, with extensive crosstalk among them that warrants further exploration.

In the photoperiod pathway, the circadian clock integrates signals from the length of the day to regulate the GIGANTEA (GI)-CONSTANS (CO)-FLOWERING LOCUS T (FT) module (Imaizumi et al., 2005; Sawa et al., 2007; Takagi et al., 2023). The circadian clock orchestrates the rhythmic expression of *GIGANTEA* (*GI*), peaking before dusk. During long days (LD), light induces GI to form a complex with FLAVIN-BINDING, KELCH REPEAT, F BOX 1 (FKF1) E3 ubiquitin ligase before dusk, leading to the degradation of CYCLING DOF FACTOR1 and its homologs (CDFs), which otherwise repress *CO* transcription (Sawa et al., 2007). In LD conditions, CO protein accumulates and interacts with various transcription factors (TFs), including B and C subunits of Nuclear Factor Y, namely NF-YBs and NFYCs, to activate *FT* expression in leaves (Kumimoto et al., 2008; 2010; Cao et al., 2014). FT, functioning as a mobile florigen, moves to the shoot apical meristem (SAM), where it initiates floral transition by promoting the expression of the SAM-expressed floral integrator *SUPPRESSOR OF OVEREXPRESSION OF CO 1* (*SOC1*) and activating *LEAFY* (*LFY*) and *APETALA1* (*AP1*) (Jaeger and Wigge, 2007). Conversely, under short days (SD), the formation of the GI-FKF1 complex is reduced due to the unsynchronized accumulation of GI and FKF1 proteins, and *CO* repression by CDFs delays flowering (Sawa et al., 2007).

While extensive research has been conducted on regulating flowering time under inductive photoperiods, the mechanisms governing the repression of floral initiation in non-inductive day lengths remain largely unexplored. The GA pathway plays a significant role in suppressing the expression of *FT* and *SOC1* under SD by orchestrating multiple flowering pathways (Bao et al., 2019). GA repressor DELLA proteins interact with and stabilize the MYC3 transcription factor, which directly represses *FT* or indirectly inhibits the association of the CO/NF-YB activator with the *FT* promoter (Cao et al., 2014; Bao et al., 2019). Furthermore, GA regulates the miR156-SQUAMOSA PROMOTER BINDING PROTEIN-LIKE (SPL) module in the age flowering pathway (Huijser and Schmid, 2011). In juvenile plants, high levels of *miR156* repress *SPL9*, *SPL10*, and *SPL15*, while decreasing *miR156* levels during maturation enhances the expression of *SPL9* and its paralogs (Wang et al., 2009). SPL9 promotes the transcription of *pri-miRNA172*, which represses AP2-like TFs, including *SCHLAFMUTZE* (*SMZ*) and *EAT1 TARGETs* (*TOEs*), direct or indirect repressors of *FT* or *SOC1* (Mathieu et al., 2009). Under SD, DELLA can interact with SPL9 to inhibit its activity, leading to delayed flowering (Yu et al., 2012). Moreover, DELLA interacts with and stabilizes FLOWERING LOCUS C (FLC), the primary floral repressor in the vernalization and autonomous pathways. FLC, along with other MADS TFs like SHORT VEGETATIVE PHASE (SVP), FLOWERING LOCUS M (FLM), and its homologs, MADS AFFECTING FLOWERINGs (MAFs), recruit histone modifiers to establish repressive epigenetic marks on the *FT* and *SOC1* loci (Gu et al., 2013; Li et al., 2016), thereby facilitating the repression of *FT* and *SOC1* by FLC.

Warm temperatures can alleviate the repression of flowering under SD through crosstalk among flowering pathways (Fernández et al., 2016; Jin and Ahn, 2021). The bHLH transcription factors PHYTOCHROME INTERACTING FACTOR 4 and 5 (PIF4 and PIF5) interact with CO to promote *FT* expression at high ambient temperatures under SD (Kumar et al., 2012; Fernández et al., 2016). The expression and activity of PIF4 can be enhanced by the clock/photoperiod regulators EVENING COMPLEX under high-temperature conditions (Silva et al., 2020). Additionally, the accessibility of PIF4 to the *FT* promoter can be regulated by the RGA DELLA protein, which interacts with the DNA-binding domain of PIF4; however, the flowering regulation by GA appears to be partially independent of PIF4 (de Lucas et al., 2008). Moreover, SVP and FLM exert more pronounced repression of *FT* and *SOC1* expression under SD conditions (Scortecci et al., 2001; Posé et al., 2013). While it has been documented that high temperatures decrease the stability of the SVP protein, thereby diminishing its repressive effects on flowering (Lee et al., 2013), the mechanism underlying the day-length-specific regulation of SVP-FLM functions remains elusive.

RNA splicing emerges as a significant contributor to the modulation of florigen and upstream regulators in response to environmental fluctuations (Posé et al., 2013; Shang et al., 2017; Park et al., 2019; Qi et al., 2019). It generates protein isoforms from spliced RNA variants; some play antagonistic roles in floral pathways, while others modulate RNA stabilities. In the thermosensory flowering pathway, low and high temperatures modulate the alternative splicing of *FLM* (Posé et al., 2013). The major FLM isoform, FLMβ, interacts with SVP to repress *FT* and *SOC1*. Under high ambient temperatures, a decrease in *FLMβ* and an increase in *FLMδ*, which leads to FLMδ protein competing with FLMβ for binding to SVP and then derepressing of *FT* and *SOC1*. In a similar scenario to the photoperiod pathway, *CO* produces two isoforms in a ratio determined by the photoperiods (Gil et al., 2017). COα promotes the expression of *FT*, whereas COβ, encoding a truncated CO protein, interacts and negatively regulates COα by enhancing its degradation by COP1. The ratios of COβ to COα are higher in SD compared to LD. Furthermore, *PIF4*, another ambient-temperature-sensing flowering regulator, transcribes alternative isoforms (Huang et al., 2022). PIF4-I4R (with 4th intron retained), identified as a translatable splicing isoform, produces a protein that can interact with full-length PIF4, though its specific function remains unknown. The significance of RNA alternative splicing in floral regulation is underscored by observations that mutations in RNA splicing machinery components often lead to altered flowering phenotypes (Shang et al., 2017; Park et al., 2019).

The RNA splicing is processed by spliceosomes, and their activities are dynamically regulated by splicing factors associated with them. One such factor is the MOS4-ASSOCIATED COMPLEX (MAC complex), an evolutionarily conserved splicing factor that is crucial for plant RNA splicing (Monaghan et al., 2009; Jia et al., 2017; Li et al., 2019). In Arabidopsis, the MAC complex consists of MAC3A and MAC3B, a pair of U-box type E3 ubiquitin ligase homologs, whose C-terminal WD40 domains predicted to interact with MOS4, AtCDC5, and PRL1 as MAC core proteins. The E3 ubiquitin ligase functions lie in the N-terminal U-box domains of MAC3A and MAC3B to interact with E2 ubiquitin-conjugation enzyme (E2) and facilitate ubiquitin transfer to their substrates. In human cells, the homolog of MAC3A and MAC3B, PRP19, not only transfers ubiquitin to itself for auto-inhibition, it can ubiquitinate the K63 of PRP3, a specific subunit of U4 snRNP, to enhance the association of the U4/U6/U5 snRNP within the spliceosome (Song et al., 2010; de Moura et al., 2018). While the E3 ubiquitin ligase functions of MAC3A and MAC3B in plants remain to be fully explored, their regulatory roles in plant RNA splicing have been evidenced by transcriptome analysis, which revealed misregulation of approximately 11,000 genes in 9-day-old *mac3a/mac3b* double mutant seedlings (Li et al., 2019). These splicing defects may explain a wide range of phenotypes, including circadian rhythm lengthening, growth abnormalities, altered responses to abiotic stress, changes in defense mechanisms, and defects in miRNA biogenesis (Monaghan et al., 2009; Jia et al., 2017; Tu and Chen, 2022; Jiang et al., 2023). Previous reports on the flowering phenotypes under LD conditions have shown that *mac3a* and *mac3b* single knockout mutants, or the *mac3a/mac3b* double knockdown mutant (the double null mutant being embryonic lethal), exhibit either no differences or moderate delays in flowering (Li et al., 2019; Feke et al., 2020).

Given the pivotal role of alternative splicing in regulating floral transitions and the significant alterations of splicing events observed in the *mac3a/mac3b* Arabidopsis seedlings, the previously reported phenotypes under long-day (LD) conditions cannot fully account for this discrepancy (Jia et al., 2017; Li et al., 2019; Feke et al., 2020). Here, we present evidence demonstrating that the Arabidopsis *mac3a/mac3b* double mutant exhibits significantly early flowering under SD while displaying a phenotype similar to the wild type under LD conditions. RNA-sequencing analyses of SD-grown *mac3a/mac3b* mature plants revealed expression and splicing profiles of floral regulators in multiple flowering pathways that were markedly altered. One potential cause for the early flowering phenotype is the involvement of *FLM*, a floral regulator in the ambient temperature flowering pathway. MAC3A/MAC3B can associate with *FLM* RNA to regulate the ratios of *FLMβ* and *FLMδ*, thereby influencing flowering repression. The daylength-specific transcription of *MAC3A* and *MAC3B* may associated with their regulatory role in flowering only under SD. These findings suggest that the MAC complex might be a central hub coordinating crosstalk among the flowering pathways by regulating RNA splicing under short-day conditions.

## Results

### *MAC3A* and *MAC3B* repress flowering in short-day

To comprehensively investigate the roles of *MAC3A* and *MAC3B* in flowering regulation, we measured the flowering times under both long-day (LD, 16-hour-light/8-hour-dark) and short-day (SD, 8-hour-light/16-hour-dark) conditions (**Figure 1a-b; Supplemental Table s1**). The single mutants of *mac3a* and *mac3b*, as well as the *mac3a/mac3b* double mutant, displayed flowering times similar to the wild type under LD, as determined by the number of rosette leaves at bolting (**Figure 1a, left panel**), aligning with earlier findings by Li et al. (2019). When the flowering time was assessed by the day of bolting, the *mac3a/mac3b* double mutant manifested a minor delay, echoing the observations of Feke et al. (2020) (**Figure 1a, right panel**). Notably, under SD conditions, the *mac3a/mac3b* double mutant exhibited significantly earlier flowering based on both flowering measurement indices, opposite to phenotype in LD (**Figure 1a**). Further analyses of the floral integrator transcripts, *FT* and *SOC1*, showed elevated peak expressions in the *mac3a/mac3b* double mutant compared to Col-0 under SD (**Figure 1c**). This finding supports early flowering phenotypes in the *mac3a/mac3b* mutant under SD.

**Figure 1.**
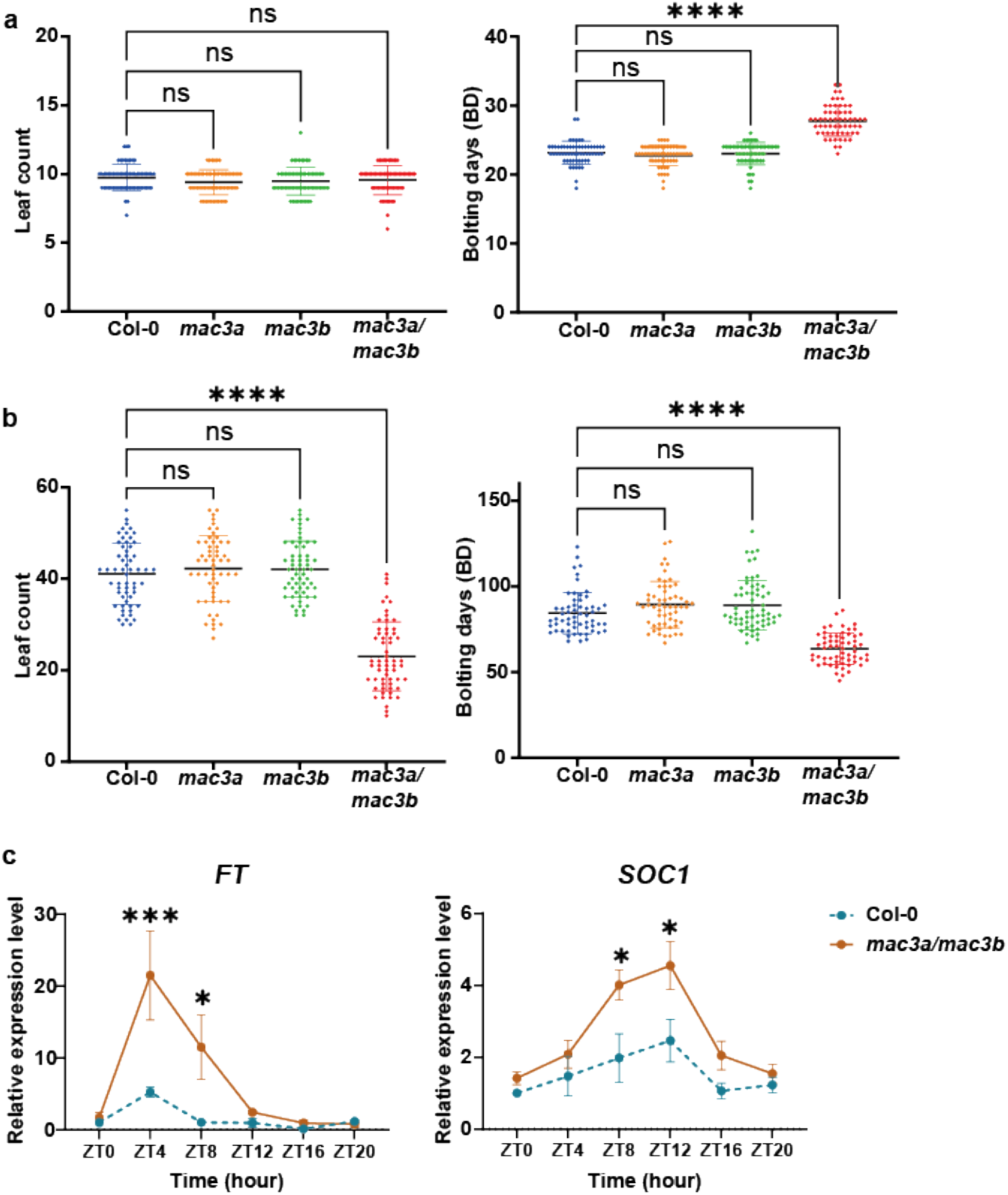
The flowering phenotypes of Col-0, *mac3a*, *mac3b*, and *mac3a/mac3b* mutants in long-day (LD) and short-day (SD) The flowering time of Col-0, *mac3a*, *mac3b*, and *mac3a/mac3b* mutants grown under **(a)** LD (16h light/8h dark) or **(b)** SD (8h light/16h dark) was measured by rosette leaf number (left panel) when inflorescence reached 7 cm and bolting day (right panel) when inflorescence reached 1 cm. Data were collected from three biological replicates (n≥ 56) and analyzed using ANOVA and Kruskal-Wallis test: ****, p< 0.0001; ns, not significant. The numeric data can be found in Supplemental Table s1. **(c)** The transcript levels of *FT* (left panel) and *SOC1* (right panel) of Col-0 and *mac3a/mac3b* seedlings in SD. The samples were harvested every 4 hours after light on (Zeitgeber time 0, ZT0), and the expression levels were measured by RT-qPCR. The relative expression levels were normalized with *UBQ10* and compared to the Col-0 at ZT0. The error bars indicated standard deviations from three biological replicates. Data were analyzed using two-way ANOVA and Šídák’s test: ***, p< 0.001; *, p< 0.05.

### The flowering phenotype of *mac3a/mac3b* mutant does not correlate with *CONSTANS* levels

To ascertain if *CO* mediated the upregulation of *FT* and *SOC1* in the *mac3a/mac3b* mutant, we analyzed the expression patterns of *CO*. The *CO* expression in the *mac3a* and *mac3b* single mutants or *mac3a/mac3b* double mutant was comparable to Col-0 in SD or LD, respectively (**Figure 2a**). Remarkably, further examination of the diurnal expression profiles of *CO* in the *mac3a/mac3b* double mutant compared to wildtype under SD conditions exhibited similar patterns, which did not account for the observed flowering phenotypes nor the expression of *FT* (**Figures 2b and 1b-c**). Given the roles of MAC3A and MAC3B in RNA splicing, we next investigated whether these factors might influence the balance between two splice variants of *CO*, *COα* and *COβ*, which in turn could modulate *FT* expression (**Figure 2c-d; Supplemental Figure s1**). However, the *mac3a/mac3b* mutant presented a reduced *COα* to *COβ* ratio due to increased *COβ* expression, theoretically suppressing *FT* expression. This pattern cannot explain the elevated *FT* expression nor the early flowering observed in the *mac3a/mac3b* mutant in SD.

**Figure 2.**
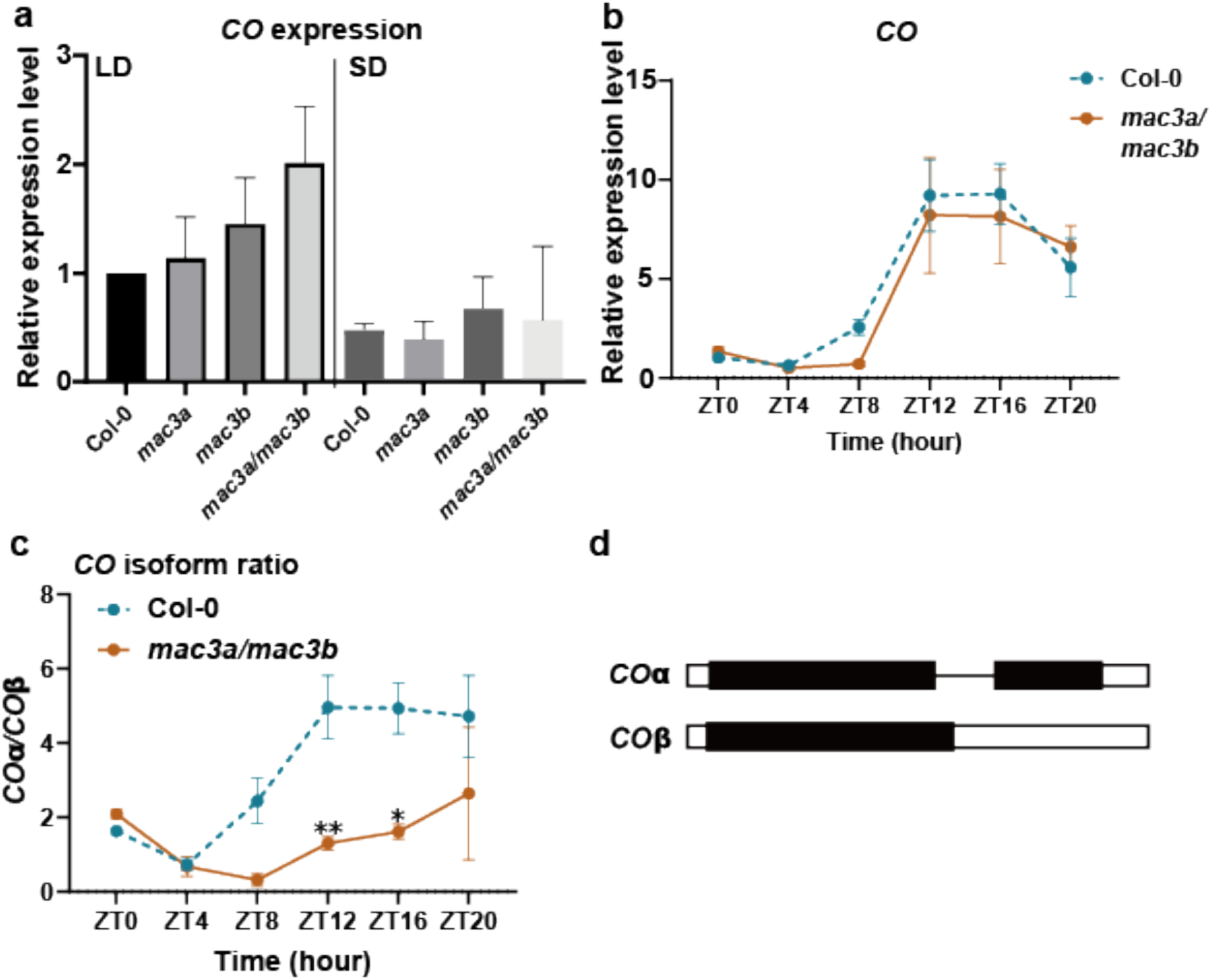
The expression of *CO* and its isoforms measured by RT-qPCR. **(a)** The relative transcript levels of *CO* in Col-0, *mac3a*, *mac3b*, and *mac3a/mac3b* mutants harvested at ZT16 under LD (16h light/8h dark) and ZT8 under SD (8h light/16h dark). **(b)** The relative diurnal transcript levels of *CO* in Col-0 and *mac3a/mac3b* mutants. The samples were harvested every 4 hours since dawn (ZT0). All time points were normalized with *UBQ10* expression and compared to the Col-0 sample at ZT0. **(c)** The ratios of *COα* over *COβ* in Col-0 and *mac3a/mac3b* mutants in SD. The expression of *COα* and *COβ* in Col-0 and *mac3a/mac3b* mutants can be found in Supplemental Figure s1. The error bars indicated standard deviations from three biological replicates. Data were analyzed using two-way ANOVA and the Šídák’s test: **, p< 0.01; *, p< 0.05. **(d)** The diagram of transcript structure and the resulting protein isoforms of *COα* and *COβ*. The rectangles are exons, with the black-filled regions representing the protein-coding regions, and the line connecting them is the intron.

### Transcriptome analyses of *mac3a/mac3b* mutant indicate alterations in various flowering pathways

Recognizing that MAC3A and MAC3B can impact transcriptome by supporting proper RNA splicing, we posited that their regulatory function in splicing might underlie the accelerated flowering observed in *mac3a/mac3b* mutants under SD conditions. To test this hypothesis, we performed RNA sequencing on 40-day-old SD-grown *mac3a/mac3b* mutants and wild-type plants, which were still in their vegetative growth phase. The analysis revealed 605 differentially expressed genes (DEGs) between the *mac3a/mac3b* mutants and Col-0 (|log_2_(fold change)|≥ 1.5 and FDR≤ 10%) (**Figure 3a; Supplemental Table s2**). Of these, 371 genes were upregulated, while 234 were downregulated. Gene Ontology (GO) enrichment analysis of the DEGs highlighted significant associations with biological processes such as circadian rhythm, response to stimulus, and post-embryonic development (**Figure 3b; Supplemental Table s3**).

**Figure 3.**
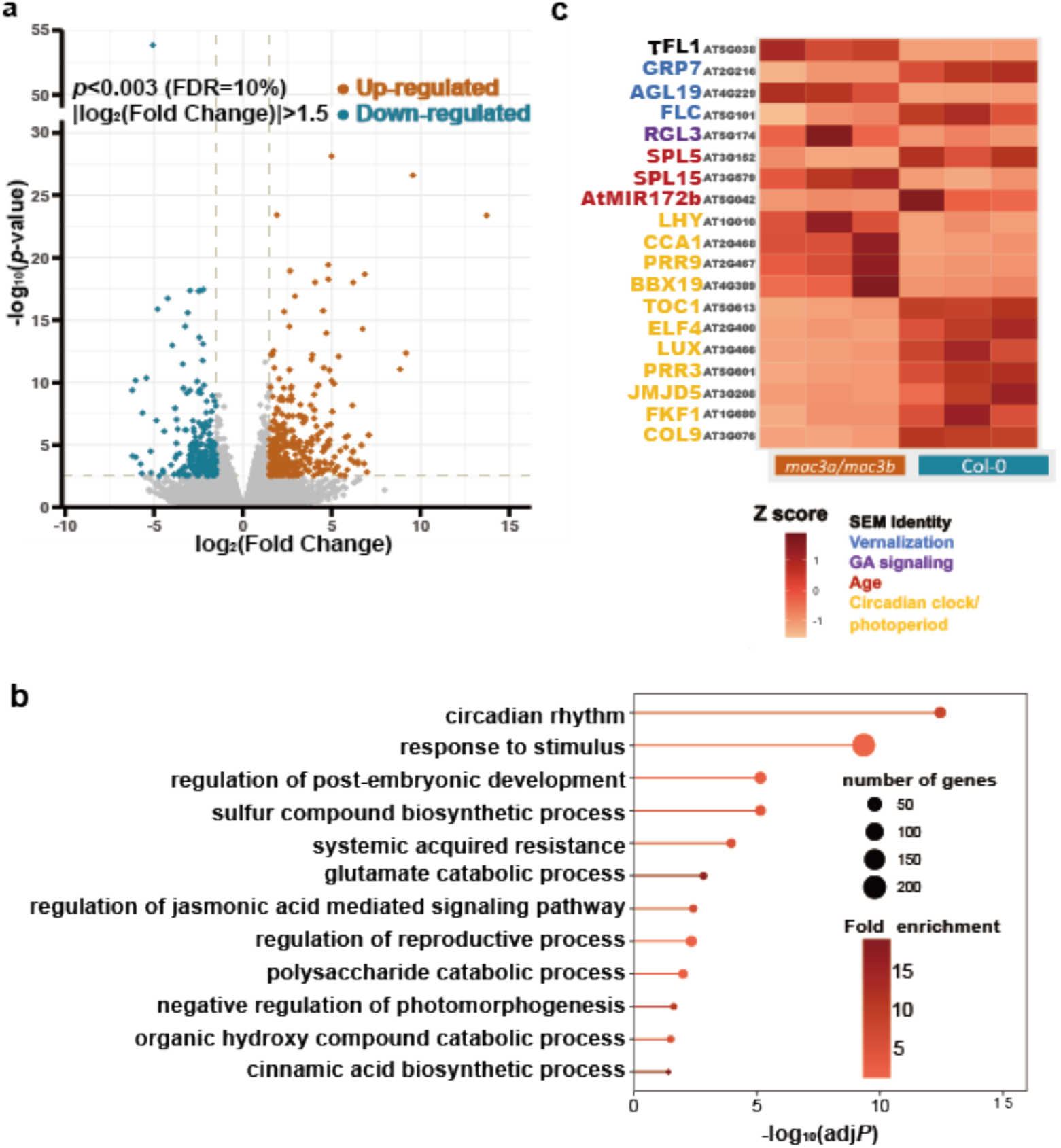
The differentially expressed genes (DEG) of SD-grown *mac3a/mac3b* mutant and their Gene ontology (GO) analysis. **(a)** The volcano plot of differentially expressed genes (DEG) in *mac3a/mac3b* mutants under SD compared to that in Col-0. The fold changes were calculated by comparing the means of normalized reads in three biological replicates of *mac3a/mac3b* and Col-0, and the p-values were calculated by T-Test of normalized reads. The up-regulated genes were marked in brown-orange, and the down-regulated genes were marked in blue with the criteria of |log_2_(fold change)|> 1.67 and 10% FDR of accepted P values. The complete gene list is in Supplemental Table s2. **(b)** The top-ranked gene ontology (GO) of the biological processes category for the DEG. Selected GO terms were plotted on the lollipop plot, with gene numbers for each GO term in the DEG list represented by the dot size, and the darkness of the lollipops indicated the fold enrichment. The list of all GO terms was included in Supplemental Table s3. **(c)** Heat map of DEG genes involved in flowering regulation. The color represents the Z-score, calculated by the normalized gene expression reads between samples. The genes belonging to specific flowering pathways were color-coded. FDR: false discovery rate.

To explore the affected flowering pathways in the *mac3a/mac3b*, we compared the DEG list with genes involving flowering initiation classified by Bouché et al. (2016) (**Supplemental Table s4**). It revealed DEG across multiple flowering pathways, including circadian clock/photoperiod, vernalization, age, and GA pathways in the *mac3a/mac3b* mutants (**Figure 3c**). Notably, morning-phased central clock regulators, like *CCA1*, *LHY*, and *PRR9,* were upregulated, whereas evening-phased regulators, such as *ELF4, LUX, TOC1*, and *PRR3,* exhibited reduced expression. This shift suggests a circadian clock phase delay, aligning with the report of lengthening the clock period in the *mac3a/mac3b* mutants by Feke et al. (2019). This result further implies that early flowering in the *mac3a/mac3b* mutants is regulated through the CO-independent pathway.

Investigating beyond photoperiodic influences, we found that the age and vernalization pathways also experienced significant alterations (**Figure 3c**). In the age pathway, the *miRNA156*-targeted flowering positive regulators, *SPL5* and *SPL15*, were downregulated and upregulated, respectively. Furthermore, SPL15-regulated pri-*miRNA172*, which acts as a floral activator, showed decreased transcript levels. These changes did not align with the same directions of early flowering observed in the *mac3a/mac3b* mutants, suggesting the age pathway might not primarily drive the phenotype. Whereas in the vernalization pathway, the floral repressor *FLC* was downregulated, and the floral activator *AGL19* was upregulated. With further confirmation by RT-qPCR, the *mac3a/mac3b* mutants exhibited overall lower *FLC* expression compared to the wildtype under SD, and the other two MADS TFs, FLM, and SVP, which complex with FLC to repress *FT*, were not significantly changed (**Supplemental Figure s2a-c**). These data suggests that *MAC3A/MAC3B* partially modulates the flowering time under SD conditions through *FLC* expression levels in the vernalization/autonomous pathway.

### Mutations in *MAC3A* and *MAC3B* lead to severe RNA splicing defects

Given the established involvement of *MAC3A/MAC3B* in RNA splicing, prior research has highlighted a marked increase in abnormal alternative splicing events, predominantly intron retention, in *mac3a/mac3b* mutant seedlings (Li et al., 2019). However, the intron retention events within the floral regulatory pathways remained unclear. We, therefore, investigated the alterations of RNA splicing to elucidate the genetic contributors to the early flowering observed in *mac3a/mac3b* mutants under SD conditions. Consequently, we identified differentially alternatively spliced transcripts in our RNA-sequencing dataset with 12,031 events encoded by 6,836 genes differential alternative splicing events (DASs) in the *mac3a/mac3b* mutants (**Table 1**). Consistent with earlier findings, the analysis revealed that over 98% of DASs or 11,876 events within 6,778 genes in the *mac3a/mac3b* mutants were characterized by increased intron retention (IR) (**Table 1**, **Figure 4a, Supplemental Table s5**). The remaining DASs were exon skipping (ES) and alterations in donor/acceptor splice sites (AltDA) involving transcription from 150 genes (**Figure 4b**, **Table 1, Supplemental Table s6**).

**Figure 4.**
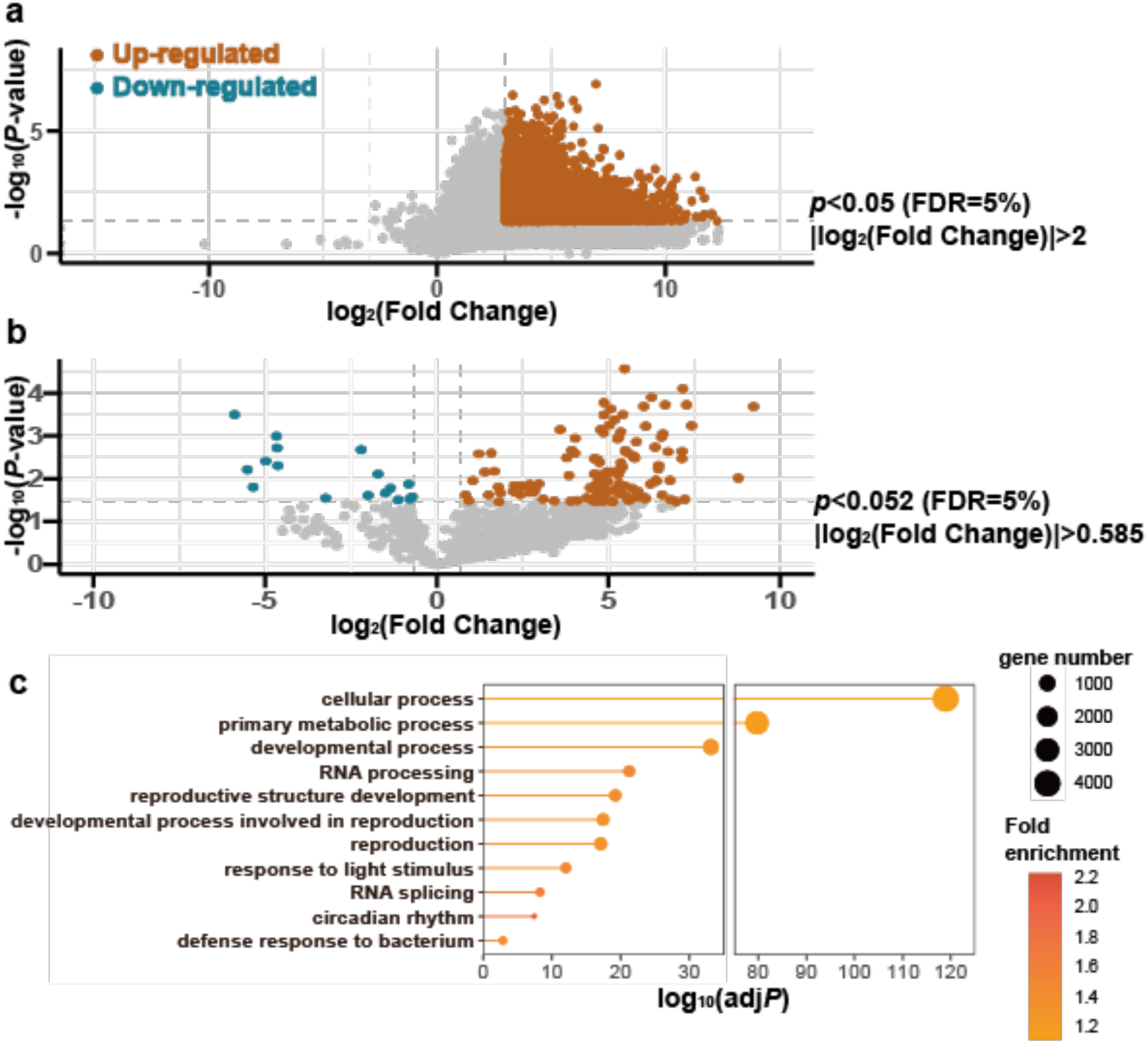
The differentially alternative spliced (DAS) events of SD-grown *mac3a/mac3b* mutant and their Gene ontology (GO) analysis. **(a)** The volcano plot of differentially intron retention (DIR) events between Col-0 and *mac3a/mac3b* identified by supporting reads ratio (|log_2_(fold change)|> 3, FDR< 0.05). **(b)** The volcano plot of differentially exon skipping (DES) events between Col-0 and *mac3a/mac3b* identified by supporting reads ratio (|log_2_(fold change)|> 0.585, FDR < 0.05). **(c)** The top-ranked gene ontology (GO) of biological processes for all differential alternative spliced events. Selected GO terms were plotted on the lollipop plot, with gene numbers exhibiting in the dot size and the darkness of the lollipops for the fold enrichment. The complete gene and GO lists were provided in Supplemental Tables s5-7. FDR: false discovery rate.

**Table 1.** The differentially alternative spliced (DAS) events in the *mac3a/mac3b* mutant compared to Col-0 under SD. The number of DAS events, including intron retention (IR), exon skipping (ES), alternative donor/acceptor splicing site (AltDA), and their corresponding gene number was listed. The IR, AltDA, or ES events were identified by the Chi-square test of reads with or without IR, AltDA, or ES in between Col-0 and *mac3a/mac3b* mutants, respectively. Out of the identified alternative splicing events, the differentially alternative splicing events were selected by significance and fold-change among replicates. The false discovery rate (FDR) of accepted p-values is limited to 5%. The list of genes and events is provided in Supplemental Tables s5 and s6.

| *mac3a/mac3b* v.s. WT |  |  |  |  |
| --- | --- | --- | --- | --- |
|  | Events | % | Genes | % |
| **total** | **12031** | **100.00** | **6836** | **100.00** |
| **IR** | **11876** | **98.71** | **6778** | **99.15** |
| **ES** | **150** | **1.24** | **146** | **2.14** |
| **AltDA** | **5** | **0.04** | **4** | **0.06** |

GO analysis of the DASs showed significant enrichment of terms associated with developmental processes and the RNA splicing machinery (**Figure 4c, Supplemental Table s7**). Additionally, the altered RNA splicing in *mac3a/mac3b* mutants affected genes integral to primary metabolic pathways, RNA processing, and responses to environmental stimuli, potentially leading to the observed pleiotropic effects on flowering initiation and growth attenuation (**Figure 1b**; Li et al., 2018; Li et al., 2019; Feke et al., 2020; Guo et al., 2024).

A subsequent comparison of genes within the DAS with established regulators of flowering initiation (**Supplemental Table s4**, Bouché et al., 2016) revealed a significant overlap: 136 genes from the DAS list were among the 306 known to regulate flowering (**Figure 5a; Supplemental Tables s4-s6**). As noted, 135 of these genes exhibited upregulated retained introns in the *mac3a/mac3b* mutant. These results underscore the critical role of *MAC3A/MAC3B* in safeguarding splicing fidelity under SD conditions.

**Figure 5.**
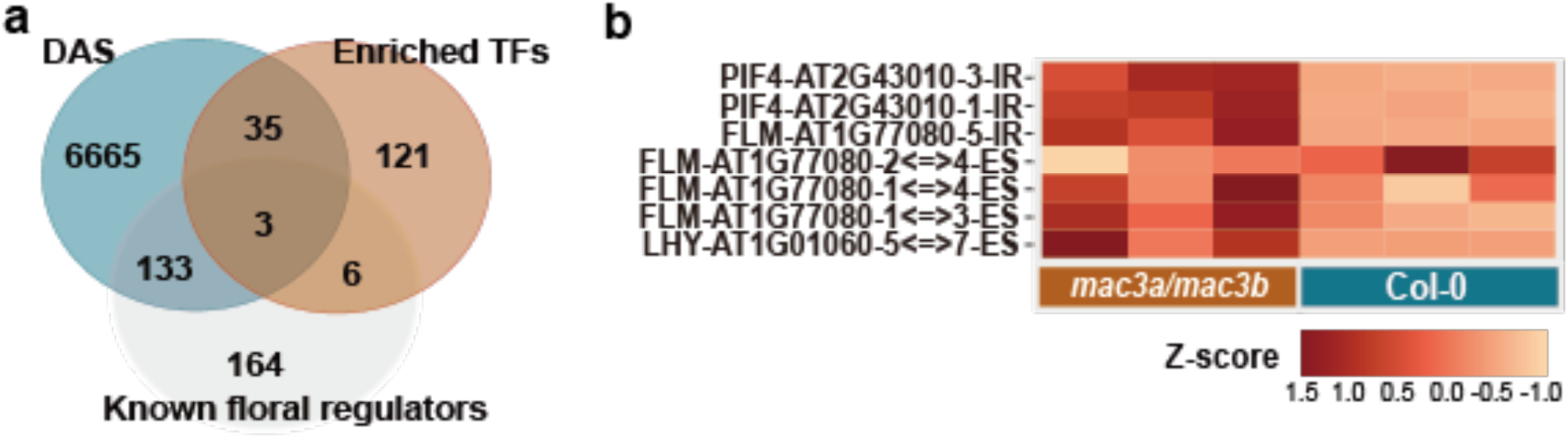
Identification of potential alternatively spliced targets of MAC3A and MAC3B for their flowering time regulation. **(a)**The Venn diagram represents the intersection of genes with differentially alternative spliced (DAS) events (Blue, Supplemental Tables s5-s6), the enriched transcription factors (TFs) upstream of the differentially expressed gene (DEG) (Orange, Supplemental Table s8), and known floral regulators (Grey, Supplemental Table s4). **(b)** Heatmap of *FLM*, *PIF4*, and *LHY* DAS events found in the *mac3a/mac3b*. The color represented the Z-score among samples from respective normalized values of splicing events. The label of each row refers to the gene name, TAIR ID, and the location of the event.

### *MAC3A/MAC3B* directly regulates splicing of the *FLOWERING LOCUS M* isoforms

To elucidate the specific regulators whose DAS defects contribute to the upregulation of *FT* and *SOC1*, and consequently the early flowering phenotype in *mac3a/mac3b* mutants, we focused on identifying overlaps among the DAS gene list (**Supplemental Tables s5-s6**), known flowering regulators (**Supplemental Table s4**), and potential transcription factors (TFs) upstream of the DEGs (Enriched TFs in **Figure 5a, Supplemental Tables s8**). The list of potential TFs was obtained by analyzing the promoter regions of each DEG for potential upstream regulators, and the enriched TFs were identified based on the frequency of downstream genes present in the DEG list (**Supplemental Table s8**). Among these TFs, 38 exhibited DAS, with three playing roles in flowering initiation: *LHY*, *PIF4*, and *FLM* (**Figure 5a**).

*LHY*, a circadian clock gene acting upstream of *CO* in the photoperiod pathway, showed increasing exon skipping (**Figure 5b, Supplemental Table s6**). However, our earlier findings have indicated that *CO*’s alterations were not a primary factor for the early flowering observed in *mac3a/mac3b* mutants. Furthermore, significant retention of the 1st and 3rd introns of *PIF4* was observed in the *mac3a/mac3b* mutants (**Figure 5b, Supplemental Figure s3a, Supplemental Table s5**). The retention of 1st intron may result in nonsense-mediated decay (NMD), while the 3rd intron potentially leads to truncated protein isoforms with missing or incomplete bHLH DNA binding domain (**Supplemental Figure s3a-b**).

Other *PIF4* alternative transcripts, though not passing our selection threshold, their products may also go through NMD. Overall, it can lead to less functional PIF4 proteins. Given PIF4’s role as a positive regulator of *FT* as Kumar et al. (2012) reported, these splicing defects could be expected to decrease *FT* expression and delay flowering.

Finally, *FLM*, a repressor in the thermosensory flowering pathway, exhibited increased inclusion of the 3rd exon and reduced inclusion of the 2nd exon in *mac3a/mac3b* mutants (**Figures 5b and 6a, Supplemental Table s6**). The alternatively splicing patterns of *FLM* lead to two major isoforms, *FLMβ* and *FLMδ*, with opposing roles in regulating flowering thermosensitivity (Posé et al., 2013). *FLMδ*, characterized by the retention of the 3rd exon and exclusion of the 2nd, mitigates *FLMβ*’s repression of *FT*. The ratios of the *FLMβ* to *FLMδ* decreases in *mac3a/mac3b* mutants under SD conditions, which could explain the upregulation of *FT* and the consequent early flowering (**Figure 6a-b, Supplemental Figure s4a**). Intriguingly, this alteration was not observed under LD conditions (**Figure 6c, Supplemental Figure s4b**), indicating that *MAC3A/MAC3B*’s regulation of *FLM* splicing is SD-specific. To ascertain if *MAC3A* and *MAC3B* directly influence *FLM* splicing, RNA-immunoprecipitation (RIP) was performed using Arabidopsis expressing FLAG-tagged *MAC3A*, revealing enrichment of *FLM* transcripts with qPCR compared to the *UBQ10* control (**Figure 6d**). These findings suggest that MAC3A and MAC3B directly modulate the splicing of *FLM*, adjusting the ratio of *FLMδ* to *FLMβ* to regulate flowering time under SD conditions.

**Figure 6.**
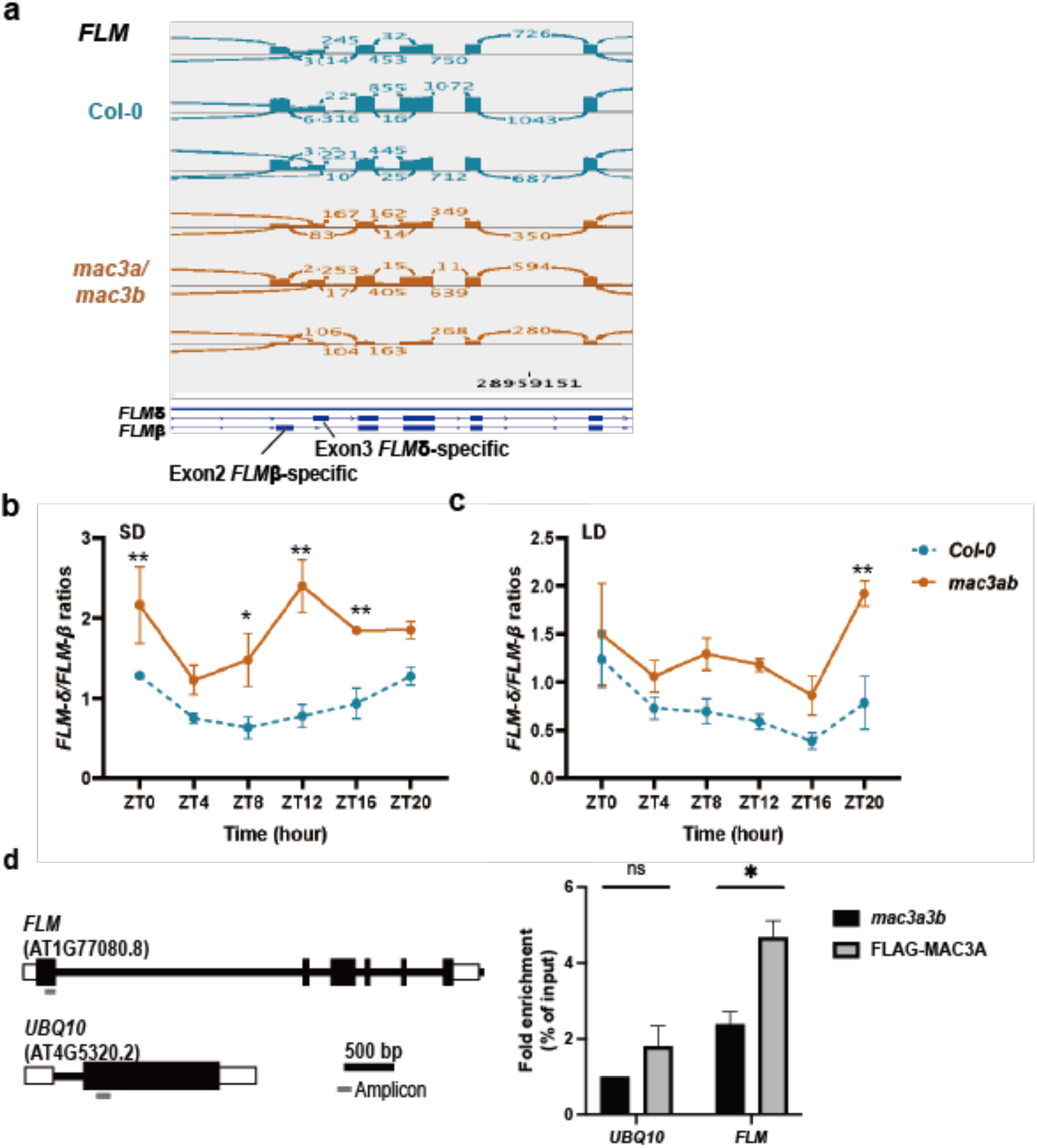
MAC3A and MAC3B directly regulate *FLM* splicing. **(a)** The sashimi plots on the *FLM* locus with highlighted exon2 and exon3 regions in Col-0 (Blue) and *mac3a/mac3b* (Orange). The height of the base represented the relative read depth, and the read number across junctions was shown between specific splicing junctions. The *FLMβ*-specific 2^nd^ exon and *FLMδ*-specific 3^rd^ exon were indicated. **(b and c)** The ratios of the *FLMδ* to *FLMβ* relative expression levels in the Col-0 and *mac3a/mac3b* in SD (b) and LD (c) were illustrated. The relative expression profiles were supplemented in Supplemental Figure s4. For each time point, the statistical analysis with a two-tailed t-test was performed (*P< 0.05 and **P< 0.01). **(d)** RNA-Immunoprecipitation followed by qPCR (RIP-qPCR) with anti-FLAG antibody in the FLAG-tagged MAC3A/*mac3amac3b* (FLAG-MAC3A) and *mac3amac3b* control. The *mac3a/mac3b* mutant and the transcript of *UBQ10* were used as control and negative control, respectively. The locations of the amplicons were illustrated in the *FLM* and *UBQ10* gene structure (left panel), and the locations of the amplicons were labeled underneath the gene structure. The enrichment of immunoprecipitated RNA was calculated relative to the input RNA and then compared to the value of *UBQ10* in the *mac3a/mac3b* mutant sample. The values shown are means ± SD. Statistical analysis was performed from three biological replicates using a one-tailed paired t-test (ns, non-significant, *P< 0.05).

### Photoperiod regulates the transcript levels of *MAC3A* and *MAC3B*

To elucidate the specific regulatory mechanisms of *MAC3A* and *MAC3B* on flowering under SD conditions, we analyzed their transcript levels under both SD and LD conditions (**Figure 7a-b**). Notably, under SD conditions, both *MAC3A* and *MAC3B* transcripts exhibited peak expression levels between 12 to 16 hours after dawn, a pattern not observed under LD conditions. These photoperiod-specific expression patterns align with the changes of *FLMδ* to *FLMβ* ratios in the under SD and LD (**Figures 4b-c, Supplemental Figure s4**). These findings indicate that photoperiod regulates *MAC3A* and *MAC3B* transcription, which affects their function in the splicing of *FLM,* potentially playing a pivotal role in modulating flowering time under SD.

**Figure 7.**
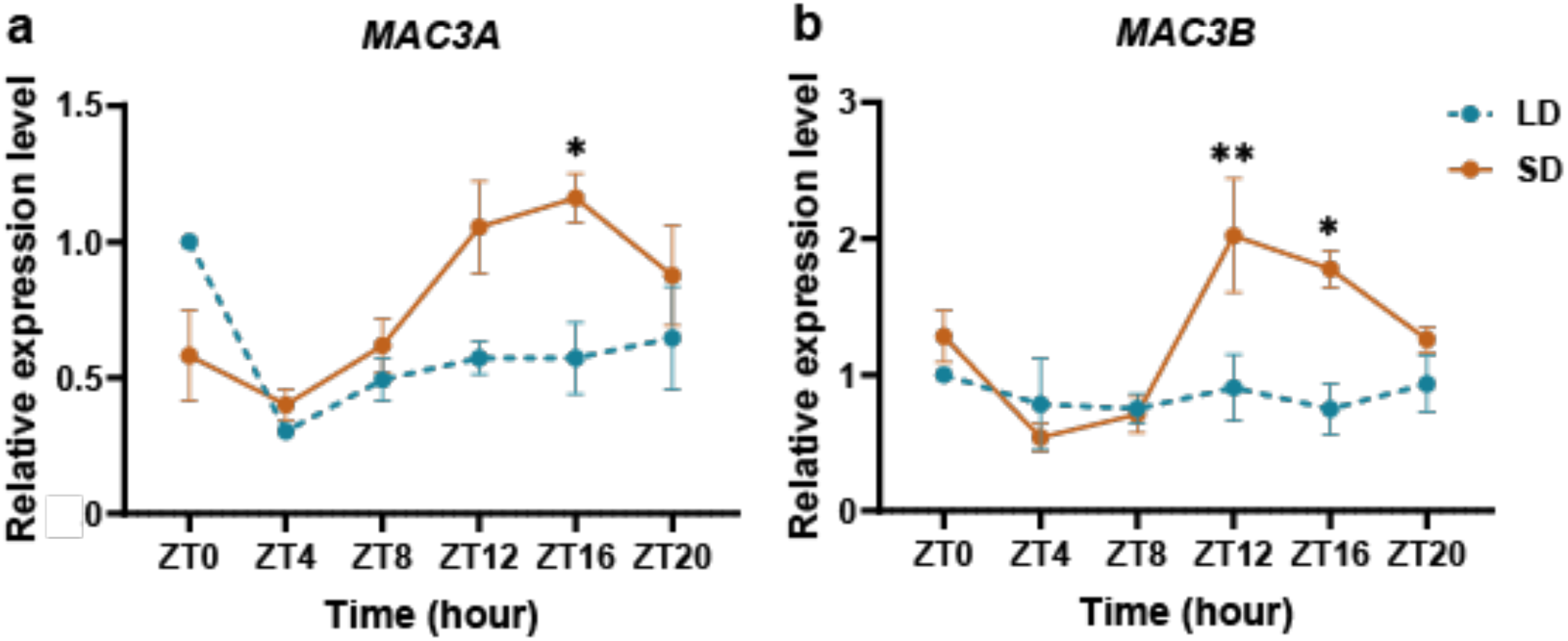
The expression of *MAC3A* and *MAC3B* in LD and SD measured by RT-qPCR. The relative diurnal expression pattern of *MAC3A* **(a)** and *MAC3B* **(b)** in LD and SD were measured with RT-qPCR. The expression was normalized with *UBQ10* and compared to the expression at ZT0 in LD. Error bars showed the standard deviations of three biological replicates, and the data were statistically tested using the two-way ANOVA and Šídák’s test: **, p< 0.01; *, p< 0.05.

## Discussion

Alternative splicing critically regulates floral transition in response to environmental cues, yet the underlying regulators and mechanisms remain largely unexplored (Shang et al., 2017; Park et al., 2019; Qi et al., 2019). Disruptions of proteins are involved in the RNA splicing process frequently result in flowering time anomalies across both LD and SD conditions. Notably, mutations in U1 snRNP components, such as *RBP45d* and *PRP39a*, or the factors associated with 3’ splicing site recognition—U2 snRNP auxiliary factors U2AF35a, U2AF35b, and U2AF65a— all lead to delayed flowering in both LD and SD (Wang and Brendel, 2006; Park et al., 2019; Chang et al., 2022). This study unveils the roles of MAC3A and MAC3B, key components of the MOS4-associated complex (MAC) splicing factor, specifically repressing floral transition under SD conditions (**Figure 8**). We discovered that the photoperiod influences *MAC3A/MAC3B* expression, driving higher expression levels in SD. While transcriptome analysis suggests that alterations in transcripts and splicing profiles across various flowering pathways contribute to regulating flowering time, our findings specifically point to MAC3A/MAC3B’s direct impact on *FLM* splicing under SD conditions. Additionally, an indirect effect on *FLC* expression emerges as another mechanism by which *MAC3A* and *MAC3B* suppress flowering under SD, highlighting a nuanced role of alternative splicing and photoperiodic regulation in floral transition. These findings also shed light on the crosstalk among floral regulatory pathways.

**Figure 8.**
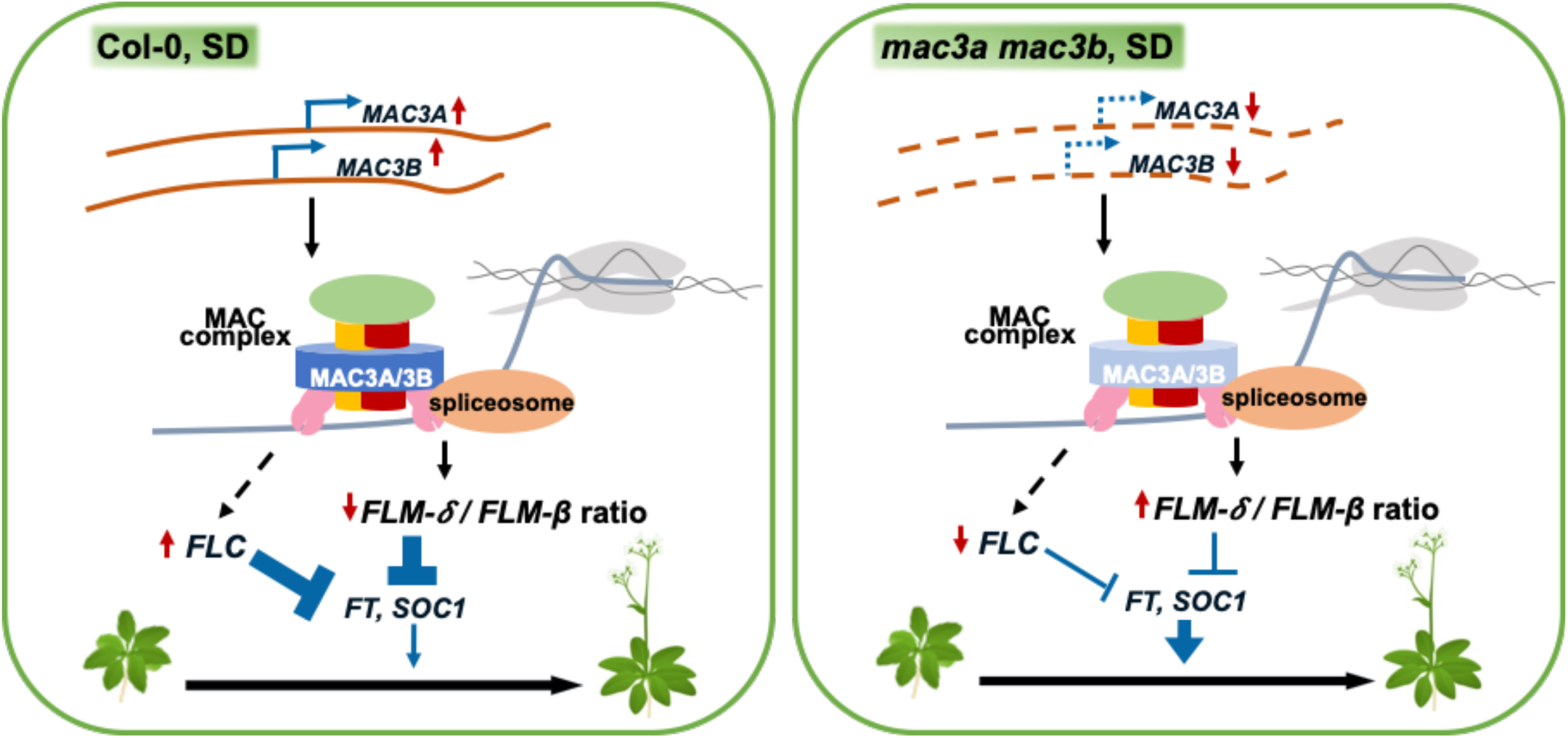
Proposed model of MAC3A and MAC3B mediated photoperiod signal to regulate flowering under SD. The working model of MAC3A and MAC3B repressing floral transition under SD. In Col-0, the expression of *MAC3A* and *MAC3B* transcription is upregulated under SD. MAC3A and MAC3B, potentially together with the MAC complex and spliceosome, facilitate proper RNA splicing of *FLM* through direct interaction. This maintains lower *FLMδ* to *FLMβ* ratios and represses floral integrators (*FT* and *SOC1*), thereby delaying the floral transition. Additionally, MAC3A and MAC3B are also required to maintain *FLC* levels to repress flowering under SD. In the *mac3a mac3b* hypomorphic mutant, reduced expression of *MAC3A/MAC3B* causes higher *FLMδ* to *FLMβ* ratios and decreased *FLC* expression under SD. Both changes increase *FT* and *SOC1* expression, resulting in early flowering.

### MAC3A and MAC3B modulate the ratios of *FLM* isoform to regulate flowering in response to photoperiod and temperature signals

Utilizing a combination of transcriptome, RIP-qPCR, and expression analysis, we have identified *FLM* as a direct splicing target of MAC3A and MAC3B under SD conditions (**Figures 5 and 6**). *FLM* functions at the crossroads of the thermosensory and autonomous flowering pathways, with its splicing of isoforms modulated by temperature (Posé et al., 2013; Jin and Ahn, 2021). The alterations of *FLM* splicing also have been found in various splicing regulator mutants under ambient temperature conditions. Our results extend this understanding by revealing that *FLM* splicing is also influenced by photoperiod, suggesting a mechanism whereby *FLM* isoform regulation serves as an integration point for photoperiod and temperature cues, facilitating flowering in response to elevated temperatures under SD conditions. This finding offers a molecular basis for previous observations of genetic interactions between *FLM-SVP* and photoperiod flowering pathways (Scortecci et al., 2001). The potential for *MAC3A* and *MAC3B* also respond to ambient temperatures to further integrate photoperiod and temperature signals through *FLM* splicing warrants additional study.

Beyond *MAC3A* and *MAC3B*, the *GLYCINE-RICH RNA-BINDING PROTEINs*, *GRP7* and *GRP8*, have been shown to regulate *FLM* isoform splicing (Steffen et al., 2019). GRP7 preferentially binds to *FLMβ* over *FLMδ* at lower temperatures under SD. The *grp7/grp8* mutants exhibit reduced *FLMβ* levels, a pattern mirrored in *mac3a/mac3b* mutants alongside decreased *GRP7* expression (**Figure 3c**), suggesting a collaborative role in modulating *FLMβ*. Future research is poised to unravel the intricate molecular and genetic interactions between *MAC3A*, *MAC3B*, and other splicing factors in adjusting *FLM* isoform ratios and orchestrating seasonal flowering responses.

### The *FLC* expression may be indirectly regulated by MAC3A and MAC3B

In SD conditions, the *mac3a/mac3b* mutants exhibit significantly earlier flowering, a phenomenon attributed, in part, to the decreased expression of the floral repressor *FLOWERING LOCUS C* (*FLC*) within the vernalization and autonomous pathways. This reduction in *FLC* expression alleviates repression on key flowering promoters *FT* and *SOC1* (**Figures 1c, Supplemental Figure s2a**). While the direct influence of MAC3A and MAC3B on *FLC* expression appears to be limited, their role in modulating *FLC* through indirect pathways is evident, particularly in the context of epigenetic regulation and post-transcriptional RNA processing (Whittaker and Dean, 2017; Qi et al., 2019). *COOLAIR*, a long non-coding antisense RNA, initially represses *FLC* during vernalization by altering the chromatin’s H3K36me3 and H3K27me3 modifications (Qi et al., 2019). In the autonomous pathway, RNA processing factors further regulate the *FLC* expression by the 3’-end processing of its antisense *COOLAIR* transcripts (Qi et al., 2019; Whittaker and Dean, 2017). Given that *MAC3A* and *MAC3B* have been established to impact miRNA biosynthesis (Li et al., 2018), their potential effect on *COOLAIR* processing and, consequently, on *FLC* expression in *mac3a/mac3b* mutants warrants further investigation.

Additionally, transcriptome analysis revealed changes in *GRP7* expression, a known interactor with *COOLAIR*, suggesting a possible regulatory link between *MAC3A*, *MAC3B*, *GRP7*, and *FLC* expression (**Figure 3c**) (Xiao et al., 2015). Interestingly, the regulatory dynamics of *MAC3A/MAC3B* and *GRP7* diverge; *MAC3A/MAC3B* seem to specifically influence *FLC* expression under SD, contrasting with *GRP7*’s broader regulatory scope across photoperiods (Xiao et al., 2015; Steffen et al., 2019). Despite *GRP7* being downregulated in *mac3a/mac3b* mutants, this does not mask the *mac3a/mac3b* early flowering phenotype (**Figures 1a, 3c**). This suggests that under SD, MAC3A and MAC3B are crucial for maintaining *FLC* expression, potentially gating the autonomous flowering pathway and highlighting a nuanced role in flowering regulation.

### Potential versatile molecular functions of MAC3A and MAC3B in regulating flowering

MAC3A and MAC3B are bona fide U-box-containing E3 ubiquitin ligases that auto-ubiquitinate themselves or other substrates (Li et al., 2018; Guo et al., 2024; Yu et al., 2024). The functions of E3 ubiquitin ligases are crucial for modulating various biological processes, including miRNA biogenesis, lateral root emergence, and organ size control. However, whether the E3 functions are required for their role in SD-specific flowering time regulation remains to be explored. In human cells, the homologs of MAC3A and MAC3B, PRP19, have been shown to ubiquitinate spliceosome subunits, affecting their assembly and activities (Song et al., 2010; de Moura et al., 2018;). This suggests that the E3 ubiquitin ligase functions of MAC3A and MAC3B might be vital for regulating splicing in the flowering pathways. Furthermore, in addition to their role as E3 ubiquitin ligases, MAC3A and MAC3B have been reported to interact with histone modifiers such as HISTONE DEACETYLASE 15 (HDA15) (Tu and Chen et al., 2022). Moreover, recent findings indicate that MAC3A and MAC3B associate with cryptochromes or other proteins to form biomolecular condensates in the nucleus under specific conditions, suggesting a potential role in regulating gene expression through epigenetic or transcriptional mechanisms beyond RNA splicing (Li et al., 2018; Jia et al., 2023).

## Materials and Methods

### Plant Materials and Growth Conditions

All *Arabidopsis thaliana* lines utilized in this study were derived from the Columbia-0 (Col-0) genetic background. The *mac3a* (SALK_089300), *mac3b* (SALK_050811), and the *mac3a/mac3b* double mutants (CS69985) were as characterized by Monaghan et al. (2009). All mutant lines were obtained from the Arabidopsis Biological Resource Center (ABRC) and confirmed through genotyping and PCR.

Seed stratification was performed by soaking in sterilized water at 4°C for three days in darkness, followed by sowing on the soil. For seedlings grown on plates, seeds underwent sterilization with 70% ethanol and 0.1% TRITON® X-100, then were plated on 1/2 strength Murashige and Skoog (MS) medium (Caisson, Cat# MSP02) supplemented with 0.8% agar (Duchefa Biochemie, Cat# P1003). Plates were kept at 4°C in the dark for three days before being moved to growth chambers. Growth conditions for plants, whether in soil or on plates, included long-day (LD) conditions (16 hours of light/8 hours of dark) or short-day (SD) conditions (8 hours of light/16 hours of dark), maintained at 22°C with light intensities ranging from 85 to 120 µmol m^-2^ s^-1^.

Flowering time was quantitatively assessed using two metrics: the date of flowering, when the inflorescence stem reached a length of 1 cm, and the total leaf count, recorded once the inflorescence stem achieved a length of 7 cm. Each experimental replicate comprised 15-20 plants per genetic line.

### Reverse Transcription Quantitative PCR (RT-qPCR)

To quantify RNA expression, *Arabidopsis thaliana* seedlings grown for 10 days under LD conditions and 14 days under SD conditions were harvested. The harvested seedlings were flash-frozen in liquid nitrogen and ground into a fine powder using a ball grinder (Retsch, MM400). Total RNA was then extracted from the tissue powder using TRIzol Reagent (Invitrogen, Cat# 15596-018) following the manufacturer’s instructions. DNase treatment (Promega, Cat# M6101) was applied to the extracted RNA samples to eliminate potential DNA contamination. Finally, the RNA concentrations and 260/280 ratios were measured with NanoDrop One (ThermoFisher) before frozen storage of the RNA samples.

For reverse transcription, 1000 ng of DNase-treated RNA was used in a 20 µL reaction employing oligo-dT primers and random hexamers according to the protocol provided by Takara (Cat# RR014B). The relative expression levels of the target genes were assessed via qPCR. For this, 4 µL of 10-fold diluted cDNA was combined with 5 µL of iQ SYBR Green Supermix (Bio-Rad, Cat# 170-8880) and 0.5 µL of each 10 µM forward and reverse primers, bringing the total reaction volume to 10 µL. These qPCR reactions were conducted using the CFX Connect Real-Time System (Bio-Rad) following a three-step PCR protocol.

This study incorporated three biological replicates for each qPCR experiment. Expression levels were normalized to the internal control gene *UBQ10* (AT4G05320). The sequences of all primers used for RT-qPCR are provided in **Supplemental Table s9**.

### RNA-sequencing Analyses of Differentially Expressed Genes (DEG) and Differential Alternative Spliced (DAS) Events

Leaves from 40-day-old Col-0 and *mac3a/mac3b* mutants grown on the soil under SD conditions were collected at Zeitgeber time 8 (ZT8). Total RNA was extracted using the RNeasy Plant Mini Kit (QIAGEN, Cat# 74904), with subsequent removal of genomic DNA contamination employing the RNase-Free DNase Set (QIAGEN, Cat# 79254). RNA-sequencing (RNA-seq) libraries for triple replicates were prepared utilizing the Illumina TruSeq toolkit (Illumina, Cat# FC-122-1001). The detailed pipelines of differential gene expression (DEG) analysis and differential alternative splicing (DAS) event detection are provided in **Supplemental Method s1**.

Gene Ontology (GO) annotations for DEGs and DAS genes (**Supplemental Tables s3 and s7**) were analyzed using the g:GOst feature on the g:Profiler website (https://biit.cs.ut.ee/gprofiler/gost; Raudvere et al., 2019) for enriched GO terms. This study used the calculation method published by Storey and Tibshirani (2003) to estimate the FDR. Data visualization was accomplished with the ggplot2 package in R (Wickham, 2016).

To identify key transcription factors (TFs) implicated in the *mac3a/mac3b* mutants (**Supplemental Table s8**), we performed TF enrichment analysis on the differentially expressed gene (DEG) lists (**Supplemental Table s2**) utilizing the FunTFBS interaction list with a cutoff p-value of <0.05 (PlantRegMap, http://plantregmap.gao-lab.org/tf_enrichment.php; Tian et al., 2020). The FunTFBS interaction list was pre-generated based on a significant Pearson correlation between binding motif frequencies and conservation scores across base pairs. TFs with at least one FunTFBS present on the differentially expressed genes were included in the TF enrichment analysis. Fisher’s exact tests were then conducted to identify TFs with significantly over-represented target genes within the input gene set, compared to their binding sites in the genomic background.

### Generation of *FLAG-MAC3A-*overexpressing Transgenic Arabidopsis

To construct *35S::6xHis-3xFLAG-MAC3A* (or called *FLAG-MAC3A*-overexpression lines) in the *mac3a/mac3b* or *mac3a* mutant background, the *MAC3A* coding sequence was subcloned into the *pB7-HFN* plant binary vector. This vector was then transformed into *Agrobacterium tumefaciens* strain *GV3101*, which was used to introduce the construct into the *mac3a/mac3b* double mutant using the floral dip method. Selection of homozygous T3 plants was performed by screening on 1/2 MS medium supplemented with 10 µg/mL glufosinate ammonium (Cyrusbiosciences, Cat# 101-77182-82-2).

### RNA-Immunoprecipitation Quantitative PCR (RIP-qPCR)

RNA immunoprecipitation (RIP) was conducted on 15-day-old *mac3a/mac3b* and MAC3A-FLAG-overexpression *Arabidopsis* seedlings harvested at ZT12, following a modified protocol from Terzi and Simpson (2009). Seedlings were cross-linked with 1% formaldehyde under vacuum infiltration for 20 minutes, with the reaction quenched using 200 mM glycine for an additional 5 minutes under vacuum. After four washes with ddH2O and drying, the samples were frozen in liquid nitrogen and pulverized.

The pulverized tissue was incubated in 30 mL of buffer 1 (400 mM sucrose, 10 mM Tris-HCl pH 8.0, 10 mM MgCl_2_, 5 mM β-mercaptoethanol, 1 mM PMSF, 1× Roche cOmplete EDTA-free Protease Inhibitor Cocktail, and 50 U/mL Invitrogen™ SUPERase•In™ RNase Inhibitor) on ice for 30 minutes. Following filtration through miracloth and centrifugation at 3000 g for 30 minutes at 4°C, the pellet was resuspended in buffer 2 (250 mM sucrose, 10 mM Tris-HCl pH 8.0, 10 mM MgCl_2_, 5 mM β-mercaptoethanol, 1 mM PMSF, 1× cOmplete EDTA-free Protease Inhibitor Cocktail, and 50 U/mL SUPERase•In™ RNase Inhibitor), then centrifuged at 12000 g for 10 minutes at 4°C. The subsequent pellet was suspended in buffer 3 (1.7 M sucrose, 10 mM Tris-HCl pH 8.0, 2 mM MgCl_2_, 5 mM β-mercaptoethanol, 1 mM PMSF, 1× cOmplete EDTA-free Protease Inhibitor Cocktail, and 50 U/mL SUPERase•In™ RNase Inhibitor), centrifuged at 16000 g for 20 minutes at 4°C, and the resulting nuclei pellets were resuspended in buffer 4 (50 mM Tris-HCl pH 8.0, 10 mM EDTA pH8.0, 1% SDS, 1 mM PMSF, 1× cOmplete EDTA-free Protease Inhibitor Cocktail, and 160 U/mL SUPERase•In™ RNase Inhibitor) for 10 minutes on ice. Nucleic acids were fragmented using a Branson Digital Sonifier SFX 250, with a cycle of 15 minutes on and 1 minute off, repeated 20 times (totaling 5 minutes), followed by centrifugation at 15000 g for 10 minutes at 4°C.

The nuclear extract was then diluted with RIP dilution buffer (16.7 mM Tris-HCl pH 8.0, 1.2 mM EDTA pH8.0, 167 mM NaCl, 0.5% Triton X-100, 1 mM PMSF, 1× cOmplete EDTA-free Protease Inhibitor Cocktail, and 160 U/mL SUPERase•In™ RNase Inhibitor) and incubated with anti-FLAG® antibody magnetic beads (Merck, Cat# M8823) overnight at 4°C with gentle rotation. Beads were washed four times in RIP wash buffer (50 mM Tris-HCl pH 8.0, 10 mM EDTA pH8.0, 300 mM NaCl, 0.5% Triton X-100, 0.1 % SDS, 1 mM PMSF, 1× cOmplete EDTA-free Protease Inhibitor Cocktail, and 40 U/mL SUPERase•In™ RNase Inhibitor) and treated with proteinase K (Sigma, Cat# p2308). RNA was extracted using TRIzol reagent (Invitrogen, Cat# 15596-018), and residual genomic DNA was removed with RQ1 RNase-free DNase (Promega, Cat# M6101). The qPCR procedures were described as RT-qPCR section with the primers listed in **Supplemental Table s9.**

### Protein Extraction and Immunoblot

Protein was extracted from *Arabidopsis thaliana* seedlings, previously ground to a powder using a freezer mill (Retsch, MM400) in liquid nitrogen. The extraction buffer contained 4 M Urea, 100 mM Tris-HCl (pH 6.8), 5% SDS, 15% Glycerol, supplemented with 0.4% protease inhibitor cocktail (Roche, Cat# 04693132001), 0.5% β-mercaptoethanol (PanReac AppliChem, Cat# A11080100), and 0.2% bromophenol blue dye (Sigma, Cat# 62625289). Samples were subsequently boiled for 5 minutes at 105°C, followed by centrifugation at 13,000 g for 5 minutes to collect the supernatant. The clarified protein extracts were either analyzed immediately or stored at -80°C.

For immunoblot analysis, around 5 μg of the protein samples were subjected to 10% SDS-PAGE for separation. Proteins were then transferred to PVDF membranes (Millipore, Cat# IPVH00010) at 4°C. The membranes were blocked with 5% non-fat milk in TBST buffer (20 mM Tris-base, 150 mM NaCl, 0.1% Tween 20) for 1 hour at room temperature, followed by overnight incubation at 4°C with a 1:5000 dilution of anti-FLAG antibody (Sigma, Cat# F1804) in the blocking solution. After washing thrice with TBST, the membranes were incubated with an HRP-conjugated secondary antibody (Promega, Cat# W4021, 1:10,000 dilution) in blocking solution, washed three times in TBST, and developed using Clarity Western ECL Substrate (Bio-Rad, Cat# 1705061) for imaging on a 5200 Chemiluminescent Imaging System (Tanon). This procedure was independently replicated with three biological samples.

### Statistical analysis

Statistical analysis for experiments was performed using the software Graphpad Prism 10. Student’s t-test, one-way ANOVA (non-parametric Kruskal-Wallis test), or two-way ANOVA were adopted. The P values of post-hoc tests used are individually indicated in the figures. For one-way ANOVA and two-way ANOVA, P values were used to indicate significant differences between multiple groups with Dunn’s or Šídák’s post-hoc statistical test, respectively (ns, no significant difference; *, P< 0.05; **, P< 0.01; ***, P< 0.001). Each experiment was performed with at least three biological replicates. The results are presented as mean values ± standard error of the mean (SE).

## List of supplemental data

**Supplemental Table s1.** The flowering time was measured by rosette leaf number and bolting day in the Col-0, *mac3a*, *mac3b*, and *mac3a/mac3b* mutants grown under LD (16h light/8h dark) or SD (8h light/16h dark).

**Supplemental Table s2.** The list of differentially expressed genes (DEG) in the *mac3a/mac3b* mutant under SD compared to Col-0.

**Supplemental Table s3.** The results of gene ontology (GO) analyses of DEG in the *mac3a/mac3b* mutant.

**Supplemental Table s4.** The list of the flowering regulators classified by Bouché et al. (2016).

**Supplemental Table s5.** The list of differentially alternative spliced (DAS) events belongs to the intron retention (IR) category in the *mac3a/mac3b* mutant under SD compared to Col-0 in SD.

**Supplemental Table s6.** The list of differentially alternative spliced (DAS) events belongs to exon skipping (DES) and alternative donor/acceptor sites (AltDA) categories in the *mac3a/mac3b* mutant under SD compared to Col-0 under SD.

**Supplemental Table s7.** The gene ontology (GO) analyses of DAS in the *mac3a/mac3b* mutant in SD.

**Supplemental Table s8.** The list of enriched transcription factors (TFs) from the promoter analysis of DEG in the *mac3a/mac3b* mutant.

**Supplemental Table s9.** List of primers used in this study.

**Supplemental Figure s1.** The expression of *CO* isoforms in SD measured by RT-qPCR

**Supplemental Figure s2.** The expression of *FLC*, *FLM*, and *SVP* in SD measured by RT-qPCR

**Supplemental Figure s3.** Alternatively spliced events of *PIF4* were identified in the *mac3a/mac3b* mutant

**Supplemental Figure s4.** The diurnal expression profiles of *FLM* isoforms in LD and SD

**Supplemental Method s1.** RNA-sequencing Analyses of Differentially Expressed Genes (DEG) and Differential Alternative Spliced (DAS) Events

## Acknowledgments

We thank Dr. Shu-Hsing Wu, Dr. Huang-Lung Tsai, Dr. Ming-Jung Liu, and Dr. Mei-Chun Cheng for valuable discussion. We also thank Shu-Jen Chou in the Genomic Technology Core Laboratory and Wen-Dar Lin at the Bioinformatics Core Laboratory of the Institute of Plant and Microbial Biology, Academia Sinica, for technical assistance. We also thank the excellent technical assistance and the facilities of Technology Commons, College of Life Science, National Taiwan University.

## Author Contributions

The study was conceptualized, and the manuscript was written by C.M.L., Y.W.H., and C.Y.T.. Y.W.H., C.Y.T., and Y.T.T. designed and executed the experiments. S.L.T., H.Y.H., and C.Y.T. analyzed the RNA-sequencing data. Y.S.W., Y.Z.C., and Y.T.L. prepared the materials and conducted the experiments.

## Data availability

The sequencing data generated in this study have been deposited into GEO with a code: GSE 261591.

## Funding

C.M.L.: NSTC-110-2311-B-002-025-, NSTC-111-2311-B-002-025-, NSTC-112-2311-B-002-025-, NTU-CC-110L893604, NTU-CC-111L893004, and NTU-CC-112L891804

S.L.T.: NSTC-111-2311-B-001-004-, NSTC-112-2311-B-001-004-

C.Y.T.: NSTC-112-2813-C-002-208-B, PhD fellowship from NSTC and NTU

Y.T.T.: NTU-CC-111L4000, NTU-CC-112L4000, NTU-CC-113L4000

Y.S.W.: MOE Industry-Academia Cooperative PhD fellowship

## References

Bao S, Hua C, Huang G, Cheng P, Gong X, Shen L, Yu H. (2019) Molecular Basis of Natural Variation in Photoperiodic Flowering Responses. Developmental Cell 50(1), 90–101.

Bouché F, Lobet G, Tocquin P, Périlleux C. (2016) FLOR-ID: an interactive database of flowering-time gene networks in Arabidopsis thaliana. Nucleic Acids Research 44(D1), D1167–1171.

Cao S, Kumimoto RW, Gnesutta N, Calogero AM, Mantovani R, Holt BF. (2014) A distal CCAAT/NUCLEAR FACTOR Y complex promotes chromatin looping at the FLOWERING LOCUS T promoter and regulates the timing of flowering in Arabidopsis. Plant Cell 26(3), 1009–1017.

Chang P, Hsieh HY, Tu SL. (2022) The U1 snRNP component RBP45d regulates temperature-responsive flowering in Arabidopsis. Plant Cell 34(2), 834–851.

de Lucas M, Davière JM, Rodríguez-Falcón M, Pontin M, Iglesias-Pedraz JM, Lorrain S, Fankhauser C, Blázquez MA, Titarenko E, Prat S. (2008) A molecular framework for light and gibberellin control of cell elongation. Nature 451(7177), 480–484.

de Moura TR, Mozaffari-Jovin S, Szabó CZK, Schmitzová J, Dybkov O, Cretu C, Kachala M, Svergun D, Urlaub H, Lührmann R, Pena V. (2018) Prp19/Pso4 Is an Autoinhibited Ubiquitin Ligase Activated by Stepwise Assembly of Three Splicing Factors. Molecular Cell 69(6), 979–992.

Feke A, Liu W, Hong J, Li MW, Lee CM, Zhou EK, Gendron JM. (2019) Decoys provide a scalable platform for the identification of plant E3 ubiquitin ligases that regulate circadian function. eLife 8, e44558.

Feke AM, Hong J, Liu W, Gendron JM. (2020) A Decoy Library Uncovers U-Box E3 Ubiquitin Ligases That Regulate Flowering Time in Arabidopsis. Genetics 215(3), 699–712.

Fernández V, Takahashi Y, Le Gourrierec J, Coupland G. (2016) Photoperiodic and thermosensory pathways interact through CONSTANS to promote flowering at high temperature under short days. Plant Journal 86(5), 426–440.

Gendron JM, Leung CC, Liu W. (2021) Energy as a seasonal signal for growth and reproduction. Current Opinion in Plant Biology 63, 102092.

Gil KE, Park MJ, Lee HJ, Park YJ, Han SH, Kwon YJ, Seo PJ, Jung JH, Park CM. (2017) Alternative splicing provides a proactive mechanism for the diurnal CONSTANS dynamics in Arabidopsis photoperiodic flowering. Plant Journal 89, 128–140.

Gu X, Le C, Wang Y, Li Z, Jiang D, Wang Y, He Y. (2013) Arabidopsis FLC clade members form flowering-repressor complexes coordinating responses to endogenous and environmental cues. Nature Communications 4, 1947.

Guo X, Zhang X, Jiang S, Qiao X, Meng B, Wang X, Wang Y, Yang K, Zhang Y, Li N Chen T, Kang Y, Yao M, Zhang X, Wang X, Zhang E, Li J, Yan D, Hu Z, Botella JR, Song CP, Li Y, Guo S. (2024) E3 ligases MAC3A and MAC3B ubiquitinate UBIQUITIN-SPECIFIC PROTEASE14 to regulate organ size in Arabidopsis. Plant Physiology 194(2), 684–697.

Huang CK, Lin WD, Wu SH. (2022) An improved repertoire of splicing variants and their potential roles in Arabidopsis photomorphogenic development. Genome Biology 23(1), 50.

Huijser P, Schmid M. (2011) The control of developmental phase transitions in plants. Development 138(19), 4117–4129.

Imaizumi T, Schultz TF, Harmon FG, Ho LA, Kay SA. (2005) FKF1 F-box protein mediates cyclic degradation of a repressor of CONSTANS in Arabidopsis. Science 309(5732), 293–297.

Jin S, Ahn JH. (2021) Regulation of flowering time by ambient temperature: repressing the repressors and activating the activators. New Phytologist 230(3), 938–942.

Jia M, Chen X, Shi X, Fang Y, Gu Y. (2023) Nuclear transport receptor KA120 regulates molecular condensation of MAC3 to coordinate plant immune activation. Cell Host & Microbe 31(10), 1685–1699.e7.

Jia T, Zhang B, You C, Zhang Y, Zeng L, Li S, Johnson KCM, Yu B, Li X, Chen X. (2017) The Arabidopsis MOS4-Associated Complex Promotes MicroRNA Biogenesis and Precursor Messenger RNA Splicing. Plant Cell 29(10):2626–2643.

Jiang B, Zhong Z, Su J, Zhu T, Yueh T, Bragasin J, Bu V, Zhou C, Lin C, Wang X. (2023) Co-condensation with photoexcited cryptochromes facilitates MAC3A to positively control hypocotyl growth in Arabidopsis. Science Advances 9(32), eadh4048.

Jaeger KE, Wigge PA. (2007) FT protein acts as a long-range signal in Arabidopsis. Current Biology 17(12), 1050–1054.

Kumar SV, Lucyshyn D, Jaeger KE, Alós E, Alvey E, Harberd NP, Wigge PA. (2012) Transcription factor PIF4 controls the thermosensory activation of flowering. Nature 484(7393), 242–245.

Kumimoto RW, Adam L, Hymus GJ, Repetti PP, Reuber TL, Marion CM, Hempel FD, Ratcliffe OJ. (2008) The Nuclear Factor Y subunits NF-YB2 and NF-YB3 play additive roles in the promotion of flowering by inductive long-day photoperiods in Arabidopsis. Planta 228(5), 709–723.

Kumimoto RW, Zhang Y, Siefers N, Holt BF 3rd. (2010) NF-YC3, NF-YC4 and NF-YC9 are required for CONSTANS-mediated, photoperiod-dependent flowering in Arabidopsis thaliana. Plant Journal 63(3), 379-391.

Lee JH, Ryu HS, Chung KS, Posé D, Kim S, Schmid M, Ahn JH. (2013) Regulation of temperature-responsive flowering by MADS-box transcription factor repressors. Science 342(6158), 628–632.

Li M, An F, Li W, Ma M, Feng Y, Zhang X, Guo H. (2016) DELLA proteins interact with FLC to repress flowering transition. Journal of Integrative Plant Biology 58(7), 642–655.

Li S, Liu K, Zhou B, Li M, Zhang S, Zeng L, Zhang C, Yu B. (2018) MAC3A and MAC3B, Two Core Subunits of the MOS4-Associated Complex, Positively Influence miRNA Biogenesis. Plant Cell 30, 481-494.

Li Y, Yang J, Shang X, Lv W, Xia C, Wang C, Feng J, Cao Y, He H, Li, Ma L. (2019) SKIP regulates environmental fitness and floral transition by forming two distinct complexes in Arabidopsis. New Phytologist 224(1), 321–335.

Mathieu J, Yant LJ, Mürdter F, Küttner F, Schmid M. (2009) Repression of flowering by the miR172 target SMZ. PLOS Biology 7(7), e1000148.

Monaghan J, Xu F, Gao M, Zhao Q, Palma K, Long C, Chen S, Zhang Y, Li X. (2009) Two Prp19-like U-box proteins in the MOS4-associated complex play redundant roles in plant innate immunity. PLOS Pathogens 5, e1000526.

Park YJ, Lee JH, Kim JY, Park CM. (2019) Alternative RNA Splicing Expands the Developmental Plasticity of Flowering Transition. Frontiers in Plant Science 10, 606.

Posé D, Verhage L, Ott F, Yant L, Mathieu J, Angenent GC, Immink RG, Schmid M. (2013) Temperature-dependent regulation of flowering by antagonistic FLM variants. Nature 503, 414–417.

Raudvere U, Kolberg L, Kuzmin I, Arak T, Adler P, Peterson H, Vilo J. (2019) g:Profiler: a web server for functional enrichment analysis and conversions of gene lists (2019 update). Nucleic Acids Research 47(W1), W191–198.

Qi HD, Lin Y, Ren QP, Wang YY, Xiong F, Wang XL. (2019) RNA Splicing of FLC Modulates the Transition to Flowering. Frontiers in Plant Science 10, 1625.

Sawa M, Nusinow DA, Kay SA, Imaizumi T. (2007) FKF1 and GIGANTEA complex formation is required for day-length measurement in Arabidopsis. Science 318(5848), 261–265.

Scortecci KC, Michaels SD, Amasino RM. (2001) Identification of a MADS-box gene, FLOWERING LOCUS M, that represses flowering. Plant Journal 26(2), 229–236.

Shang X, Cao Y, Ma L. (2017) Alternative Splicing in Plant Genes: A Means of Regulating the Environmental Fitness of Plants. International Journal of Molecular Sciences 18(2), 432.

Silva CS, Nayak A, Lai X, Hutin S, Hugouvieux V, Jung JH, López-Vidriero I, Franco-Zorrilla JM, Panigrahi KCS, Nanao MH, Wigge PA, Zubieta C. (2020) Molecular mechanisms of evening complex activity in Arabidopsis. PANS 117(12), 6901–6909.

Song EJ, Werner SL, Neubauer J, Stegmeier F, Aspden J, Rio D, Harper JW, Elledge SJ, Kirschner MW, Rape M. (2010) The Prp19 complex and the Usp4Sart3 deubiquitinating enzyme control reversible ubiquitination at the spliceosome. Genes & Development 24, 1434–1447.

Song YH, Shim JS, Kinmonth-Schultz HA, Imaizumi T. (2015) Photoperiodic flowering, time measurement mechanisms in leaves. Annual Review of Plant Biology 66, 441–464.

Steffen A, Elgner M, Staiger D. (2019) Regulation of Flowering Time by the RNA-Binding Proteins AtGRP7 and AtGRP8. Plant and Cell Physiology 60(9), 2040–2050.

Storey JD, Tibshirani R. (2003) Statistical significance for genomewide studies. PNAS 100(16), 9440–9445.

Takagi H, Hempton AK, Imaizumi T. (2023) Photoperiodic flowering in Arabidopsis: Multilayered regulatory mechanisms of CONSTANS and the florigen FLOWERING LOCUS T. Plant Communications 4(3), 100552.

Terzi LC, Simpson GG. (2009) Arabidopsis RNA immunoprecipitation. Plant Journal 59(1), 163–168.

Tian F, Yang DC, Meng YQ, Jin J, Gao G. (2020) PlantRegMap: charting functional regulatory maps in plants. Nucleic Acids Research 48(D1), D1104–1113.

Tu YT, Chen CY, Huang YS, Chang CH, Yen MR, Hsieh JA, Chen PY, Wu K. (2022) HISTONE DEACETYLASE 15 and MOS4-associated complex subunits 3A/3B coregulate intron retention of ABA-responsive genes. Plant Physiology 190(1), 882–897.

Wang BB, Brendel V. (2006) Molecular characterization and phylogeny of U2AF35 homologs in plants. Plant Physiology 140(2), 624–636.

Wang JW, Czech B, Weigel D. (2009) miR156-regulated SPL transcription factors define an endogenous flowering pathway in Arabidopsis thaliana. Cell 138(4), 738–749.

Whittaker C, Dean C. (2017) The FLC Locus: A Platform for Discoveries in Epigenetics and Adaptation. Annual Review of Cell and Development Biology 33, 555–575.

Wickham H. (2016) ggplot2: Elegant Graphics for Data Analysis. Springer-Verlag New York. ISBN 978–3-319-24277-4.

Xiao J, Li C, Xu S, Xing L, Xu Y, Chong K. (2015) JACALIN-LECTIN LIKE1 Regulates the Nuclear Accumulation of GLYCINE-RICH RNA-BINDING PROTEIN7, Influencing the RNA Processing of FLOWERING LOCUS C Antisense Transcripts and Flowering Time in Arabidopsis. Plant Physiology 169(3), 2102–2117.

Yu S, Galvão VC, Zhang YC, Horrer D, Zhang TQ, Hao YH, Feng YQ, Wang S, Schmid M, Wang JW. (2012) Gibberellin regulates the Arabidopsis floral transition through miR156-targeted SQUAMOSA promoter binding-like transcription factors. Plant Cell 24(8), 3320–3332.

Yu Z, Qu X, Lv B, Li X, Sui J, Yu Q, Ding Z. (2024) MAC3A and MAC3B mediate degradation of the transcription factor ERF13 and thus promote lateral root emergence. Plant Cell, koae047.

