## Supplemental Figures and Methods for "MAC3A and MAC3B modulate *FLM* splicing to repress photoperiod-dependent floral transition"

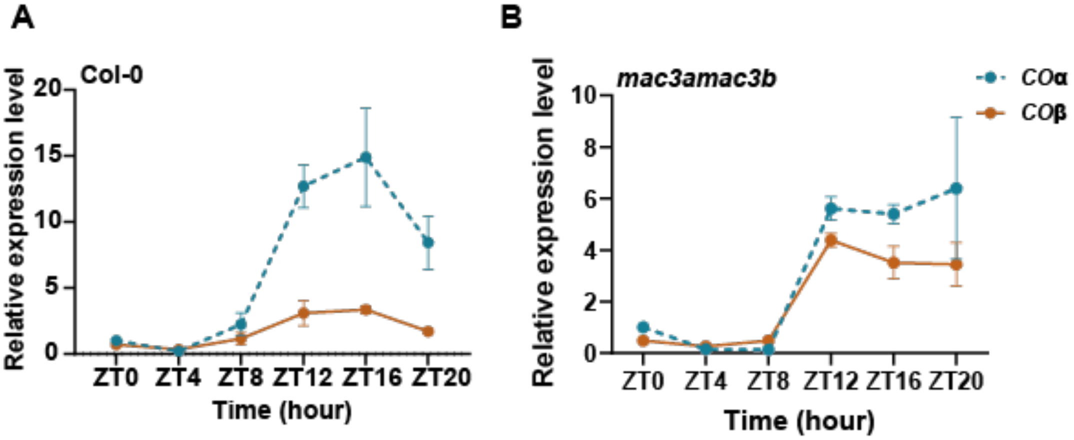


**Supplemental Figure s1. The expression of *CO* isoforms in SD measured by RT-qPCR**

The relative dirunal expression of *COα* and *COβ* in Col-0 **(a)** and *mac3a/mac3b* **(b)** in SD. The relative transcript levels of *COα* and *COβ* in Col-0 and *mac3a/mac3b* were measured from Arabidopsis grown under SD (8h light/16h dark). The samples were harvested every 4 hours since dawn (ZT0). All time points were normalized with *UBQ10* expression and compared to the *COα* at ZT0. The error bars indicated standard deviations from three biological replicates.

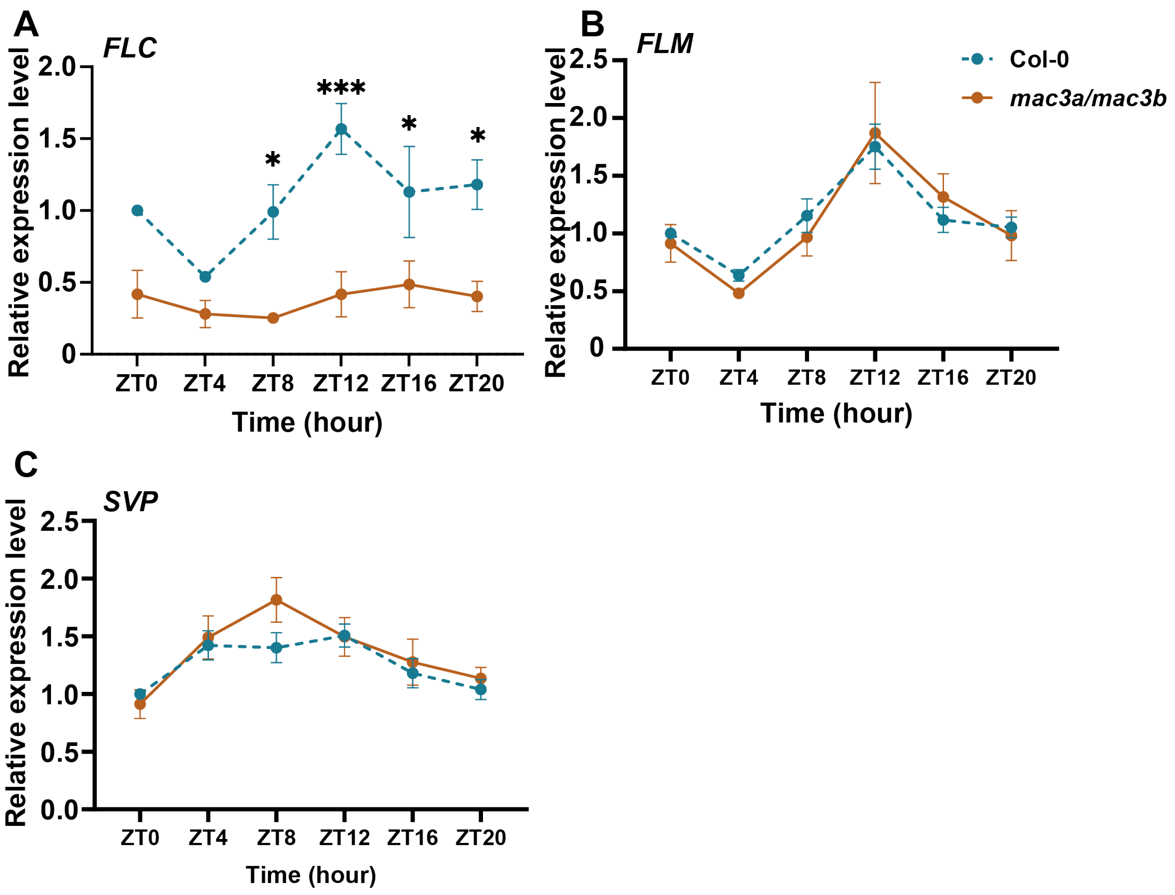


**Supplemental Figure s2. The expression of *FLC*, *FLM*, and *SVP* in SD measured by RT-qPCR**

The diurnal expression levels of **(a)** *FLC*, **(b)** *FLM*, and **(c)** *SVP* in the Col-0 and *mac3a/mac3b* mutant seedlings under SD were measured by RT-qPCR. The relative expression levels were normalized with *UBQ10* and compared to Col-0 at ZT0. Error bars indicated standard deviations of three biological replicates. Data were analyzed using two-way ANOVA and Šídák's test: ***, $p < 0.001$; *, $p < 0.05$.

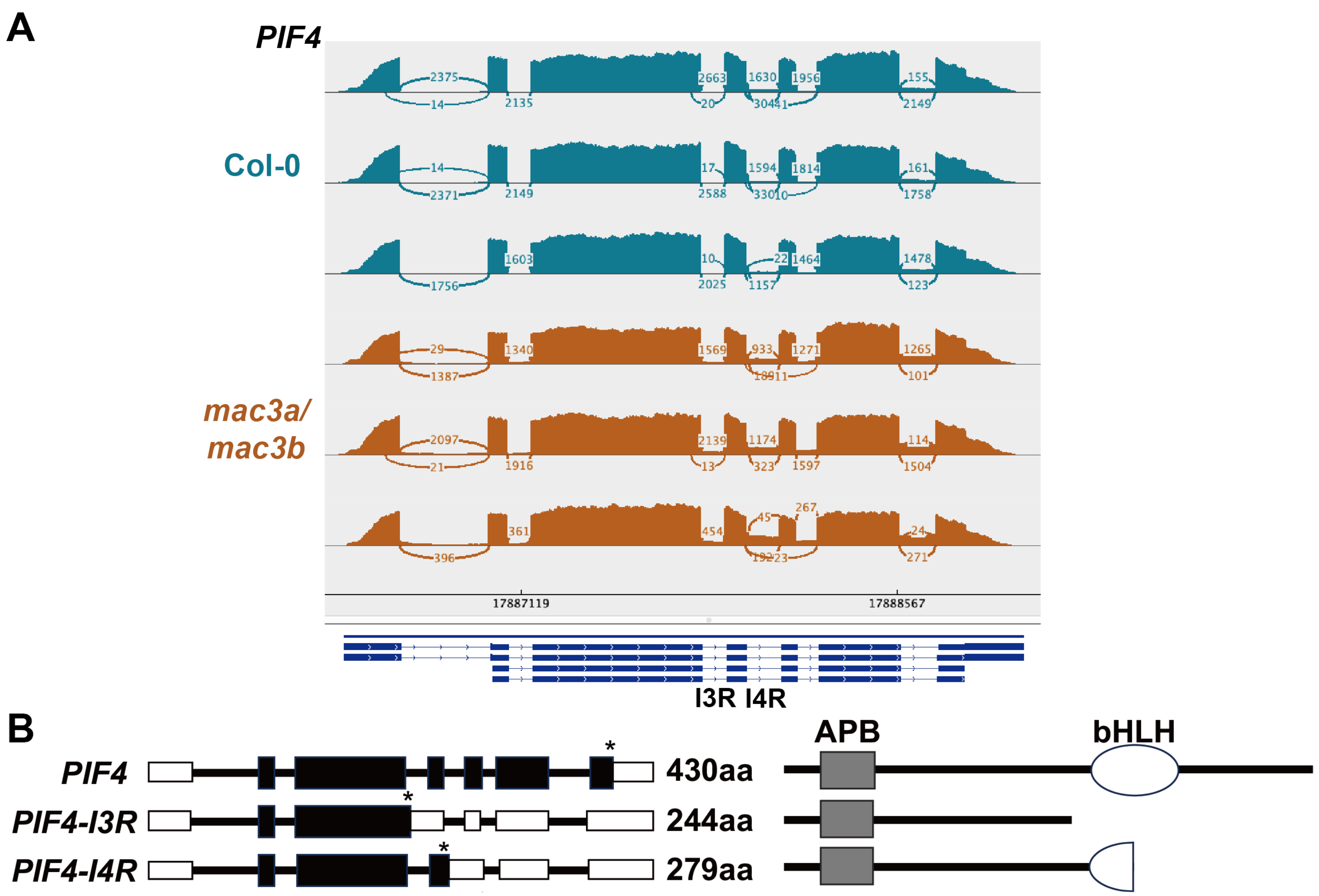


**Supplemental Figure s3. Alternatively spliced events of *PIF4* were identified in the *mac3a/mac3b* mutant**

**(a)** The sashimi plots demonstrate the splicing events found in Col-0 (Blue) and *mac3a/mac3b* (Orange) on the *PIF4* locus. The height of the base represents the relative read depth, and the read number across junctions was shown between specific splicing junctions. Intron 3, 4, and 5 (I3R, I4R, and I5R) annotated the IR events that may happen on the *PIF4* locus. In differentially alternative splicing events, only the I3R event out of these three events passes our statistical criteria. **(b)** The predicted *PIF4* isoforms with different retained introns. PIF4 and its intron-retained form of transcripts were drawn with untranslated regions (UTR) in white blocks, coding sequence (CDS) in black blocks, spliced introns in black lines, and translation stop codon in the star above (left). On the right panel, predicted peptides encoded from specific transcripts were annotated with active phyB-binding (APB, grey box) and complete/truncated bHLH (complete/sliced white oval) domains.

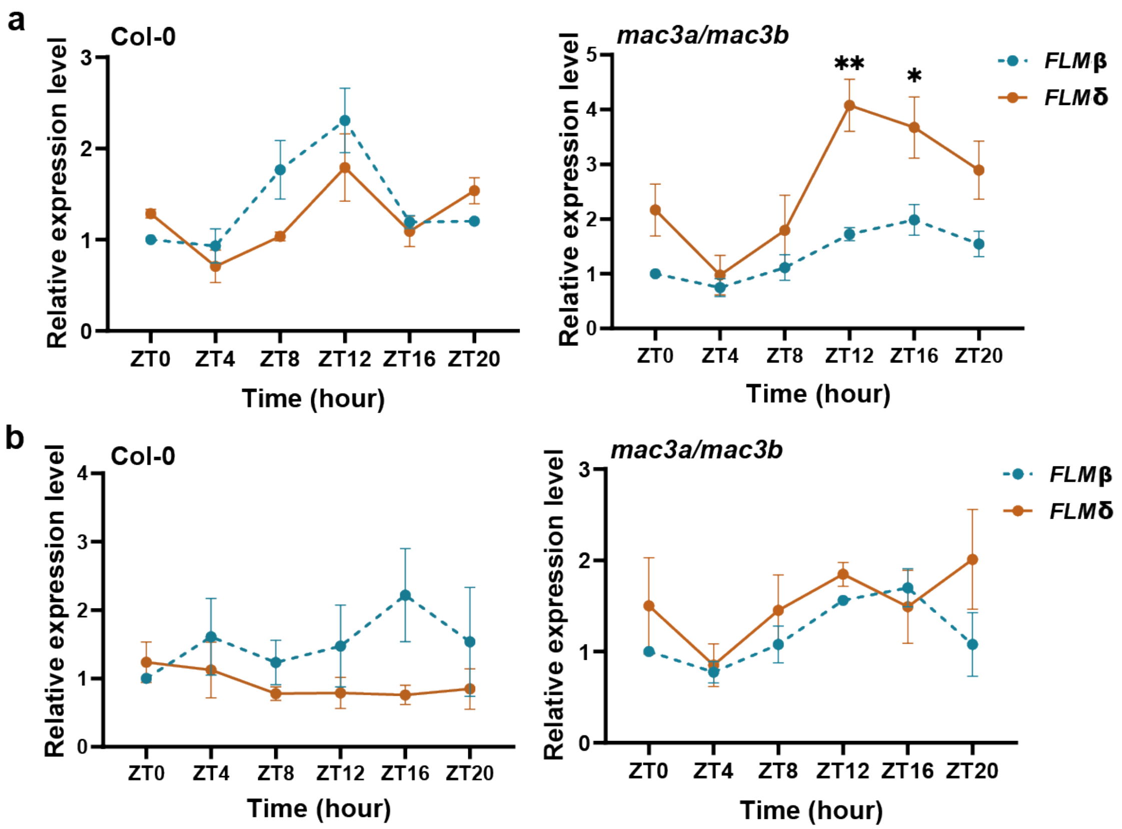


**Supplemental Figure s4. The diurnal expression profiles of *FLM* isoforms in LD and SD**

The relative expression levels of *FLMβ* and *FLMδ* were measured with RT-qPCR in Col-0 (left panel) and *mac3a/mac3b* (right panel) seedlings under SD **(a)** and LD **(b)**. All time points were normalized with *UBQ10* expression and compared to the *FLMβ* expression at ZT0. Error bars represented the standard deviations of three biological replicates. Data were analyzed using ANOVA and the Kruskal-Wallis test: *, $p < 0.05$; **, $p < 0.01$.

**Supplemental Method s1. RNA-sequencing Analyses of Differentially Expressed Genes (DEG) and Differential Alternative Spliced (DAS) Events**

The paired-end reads were first processed with Trimmomatic (Bolger et al., 2014) to excise undesired Illumina adaptor sequences and poly-G repetitions. Differential gene expression (DEG) analysis employed the STAR algorithm (Dobin et al., 2013) for alignment to the Arabidopsis TAIR10 genome using Araport11 (201606 released) annotations. Counts of aligned reads were generated with HTSeq (Anders et al., 2013), followed by normalization and comparative analysis with DESeq2 (Love et al., 2014). Statistical significance was assessed by p-values, with a false discovery rate (FDR) threshold set to 10% for identifying DEGs.

For alternative splicing event detection, reads were aligned to transcript coordinates from Araport11 using Bowtie2 (Langmead & Salzberg, 2012), applying a minimum identity threshold of 0.95. Unmapped reads were further processed with BLAT (Kent, 2002) against the TAIR10 genome, requiring an identity threshold of 0.9 and at least 8 base matches on both ends. Previously established "Read Analysis & Comparison Kit in Java" pipeline (RackJ, http://rackj.sourceforge.net) (Chang et al., 2022) then aggregated these mappings to identify sample-sensitive alternative splicing events, including intron retention, exon skipping, and donor-acceptor changes, using Chi-square tests. Events showing significant differences and fold-change among replicates were selected, setting an FDR threshold of 5%.

The IR events were identified by the Chi-square test of reads with or without IR between Col-0 and *mac3a/mac3b* mutants, with fold change criteria of $|Log_2(Fold\ change)| > 3$. Similarly, Alternative Donor/Acceptor sites (AltDA) events were identified by the Chi-square of the number of reads test with or without event-supportive exons, with fold change criteria of $|Log_2(Fold\ change)| > 1$. The statistic method used on AltDA events was applied to ES events with fold change criteria of $|Log_2(Fold\ change)| > 0.585$. Out of the identified alternative splicing events, the differentially alternative splicing events were selected by significance and fold-change among replicates. Events showing significant differences and fold-change among replicates were selected, setting an FDR threshold of 5%.

**References for the Methods**

**Anders S, Pyl PT, Huber W.** (2015) HTSeq--a Python framework to work with high-throughput sequencing data. *Bioinformatics* **31**(2): 166-169.

**Bolger AM, Lohse M, Usadel B.** (2014) Trimmomatic: a flexible trimmer for Illumina sequence data. *Bioinformatics* 2014: **30**(15): 2114-2120.

**Chang P, Hsieh HY, Tu SL. (**2002) The U1 snRNP component RBP45d regulates temperature-responsive flowering in Arabidopsis. *Plant Cell* 34(2): 834-851.

**Dobin A, Davis CA, Schlesinger F, Drenkow J, Zaleski C, Jha S, Batut P, Chaisson M, Gingeras TR.** (2013) STAR: ultrafast universal RNA-seq aligner. *Bioinformatics* **29**(1): 15-21.

**Kent WJ.** (2002) BLAT--the BLAST-like alignment tool. *Genome Research* **12**(4): 656-664.

**Langmead B, Salzberg SL.** (2012) Fast gapped-read alignment with Bowtie 2. *Nature Methods* **9**(4): 357-359.

**Love MI, Huber W, Anders S.** (2014) Moderated estimation of fold change and dispersion for RNA-seq data with DESeq2. *Genome Biology* **15**(12): 550.
